# The impact of paramutations on the invasion dynamics of transposable elements

**DOI:** 10.1101/2023.03.14.532580

**Authors:** Almorò Scarpa, Robert Kofler

## Abstract

According to the prevailing view, the trap model, invading transposable elements (TEs) are stopped in their proliferation when a TE copy jumps into a piRNA cluster, which triggers the emergence of piRNAs that silence the TE. One crucial component in the host defence are paramutations. Mediated by maternally deposited piRNAs, paramutations convert TE insertions into piRNA producing loci, thereby transforming selfish TEs into agents of the host defence. Despite this significant effect, the impact of paramutations on the dynamics of TE invasions remains unknown. To address this issue, we performed extensive forward simulations of TE invasions with piRNA clusters and paramutations. We found that paramutations significantly affect TE dynamics, by accelerating the silencing of TE invasions, reducing the number of insertions accumulating during the invasions and mitigating the fitness cost of TEs. We also demonstrate that piRNA production induced by paramutations, an epigenetically inherited trait, may be positively selected. Finally, we show that paramutations may account for three important open problems with the trap model. Firstly, paramutated TE insertions may compensate for the insufficient number of insertions in piRNA clusters observed in previous studies. Secondly, paramutations may explain the discrepancy between the observed and the expected abundance of different TE families in *D. melanogaster*. Thirdly, paramutations render piRNA clusters dispensable once the host defence has been established, which may account for the lack of TE activation when three major piRNA clusters were deleted in a previous study.

## Introduction

Transposable elements (TEs) are short stretches of DNA that selfishly multiply within genomes [41, 11]. They are highly successful, having invaded virtually all eukaryotic species investigated so far [54]. The effects of TE insertions on host fitness are still debated. While some TE insertions may confer a selective advantage [7, 13] it is likely that the majority of TEs are either neutral or deleterious to the host [35, 43, 2]. It is feasible that TEs may even threaten the persistence of species, as theoretical and experimental work suggests that TE invasions could lead to the extinction of populations [23, 53, 47]. To combat the spread of TEs, host organisms have evolved sophisticated defence mechanisms, frequently involving small RNAs [46]. In mammals and invertebrates, the host defence relies on piwi-interacting RNAs (piRNAs), small RNAs ranging in size from 23 to 29nt [15, 6]. piRNAs bound to PIWI clade proteins mediate the transcriptional and post-transcriptional silencing of TEs [6, 15, 49, 30]. These piRNAs are largely derived from discrete genomic loci, the piRNA clusters. [6]. In *D. melanogaster*, for example, piRNA clusters account for about 3% of the genome [6]. According to the prevailing view, the trap model, a TE invasion is stopped when a copy of the invading TE jumps into a piRNA cluster, which triggers the emergence of piRNAs complementary to the invading TE [60, 36, 14, 59, 42]. Several lines of evidence support this model. The insertion of artificial sequences into piRNA clusters leads to the emergence of piRNAs complementary to the inserted sequence [39, 20, 34]. A single insertion in a piRNA cluster, such as X-TAS or 42AB, is sufficient to silence a reporter [34, 19]. Finally, piRNA clusters are largely composed of many different TEs, thought to represent an ‘‘immunological memory” of past TE invasions [6, 60, 57, 56]. However, recently some doubts about the trap model have emerged. First, TE invasions in experimental populations showed that the number of cluster insertions in later generations, where the host defence has been largely established, is lower than expected under the trap model [28, 27, 47]. In agreement with this, an investigation of the composition of piRNA clusters in long-read assemblies of different *D. melanogaster* strains showed that many TE families have fewer insertions than expected [56]. Second, theoretical work shows that the number of insertions accumulating during an invasion largely depends on the size of piRNA clusters [21, 23, 51]. If the size of the piRNA clusters is known (as a percentage of the genome), quantitative predictions can be made about the expected frequency of the different TE families. However, the abundance of most families in *D. melanogaster* is much lower than expected under the trap model [23]. Third, the deletion of three major piRNA clusters in *D. melanogaster* did not lead to an activation of TEs [12]. The authors argue that dispersed piRNA-producing TE insertions outside of piRNA clusters could compensate for the loss of piRNA clusters. Such dispersed piRNA-producing loci were observed for most TE families in *D. melanogaster* [48, 56]. By mediating the installation of distinct chromatin marks, maternally deposited piRNAs are thought to direct the conversion of TE insertions into piRNA producing loci [31, 10]. This conversion of TE insertions is referred to as paramutation [10]. Paramutations were first discovered in maize and generally describe inheritable changes in the behaviour of alleles due to trans-acting (epigenetic) interactions [17]. However, apart from generating dispersed piRNA-producing TE insertions, maternally deposited piRNAs may also direct the formation of piRNA clusters in the next generation [10]. Paramutations could thus be necessary for the long-term maintenance of the location of piRNA clusters across generations (although some genomic motifs may also be involved [3]). Since paramutations depend on maternally deposited piRNAs, they can only explain the maintenance of TE silencing, but they cannot account for the emergence of novel piRNAs. One event that may trigger the emergence of such novel piRNAs is an insertion into a piRNA cluster. Therefore, piRNA clusters may be important to trigger the emergence of the first piRNAs complementary to an invading TE, but they may be dispensable once a host defence is established [9]. An alternative hypothesis suggests that siRNAs could trigger the emergence of the first piRNAs complementary to an invading TE [34]. Mediated by Dicer-2, dsRNA composed of sense and antisense transcripts of a TE may be cleaved into siRNAs that could, similarly to paramutations, turn TE-rich regions into piRNA-producing loci. Interestingly, loci that generate such siRNAs (e.g. hairpins) may not necessarily themselves be converted into piRNA-producing loci [34]. The sites that could trigger siRNA production are thus distinct from the sites that may produce piRNAs.

Previous theoretical works have studied the dynamics of TE invasions with piRNA clusters [21, 23, 51, 24, 32]. These works provided important insights. For example, they showed that, on average, about four cluster insertions are necessary to stop a TE invasion, TE invasions with piRNA clusters have three distinct phases, TE insertions in piRNA clusters may be positively selected, and piRNA clusters may prevent the extinction of populations, provided that the clusters have a minimum size [21, 23, 51, 24, 33]. Although paramutations are a crucial component of the host defence, the impact of paramutations on the dynamics of TE invasions is unclear. Investigating the effect of paramutations is especially important since it has been speculated that they may account for some of the important open problems with the trap model, such as the insufficient number of TE insertions found in piRNA clusters (see above) [56, 27]. To address this issue, we performed extensive computer simulations of TE invasions with piRNA clusters and paramutations. We also explore TE invasions under a model where siRNAs initiate the silencing of an invading TE. Using our novel simulator, InvadeGO, we show that paramutations accelerate the silencing of TE invasions, limit the amount of TEs accumulating during invasions and reduce the fitness burden of TEs. We also found that piRNA production induced by paramutations, an epigenetically inherited trait, may be positively selected. Finally, we show that paramutations can account for several of the important open issues with the trap model. First, paramutations reduce the number of cluster insertions necessary to stop a TE invasion and, therefore, may account for the low number of cluster insertions found in experimental populations. Second, since paramutations reduce the amount of TEs accumulating during TE invasion, they may explain why the TE abundance in *D. melanogaster* is lower than expected under the trap model. A model with 10% paramutable loci and 3% piRNA clusters roughly captures the abundance of the TE families in *D. melanogaster*, even in the absence of negative selection against TEs. Finally, we find that the redundancy in piRNA-producing loci generated by paramutations renders piRNA clusters dispensable once a host defence is established, explaining why Gebert et al. [12] were able to delete three major clusters without effect on TE activity.

## Results

### Model assumptions and implementation

In this work, we explore the influence of paramutations on the dynamics of TE invasions. We rely on the following definitions and model assumptions: A ‘paramutation’ refers to the conversion of a regular TE insertion - outside of piRNA clusters - into a piRNA-producing locus (fig. 1A: scenario 7). Paramutations are mediated by maternally deposited piRNAs. A ‘paramutable locus’ is a genomic region (outside of piRNA clusters) where a TE insertion could be paramutated (fig. 1A: scenarios 3, 4, 7, 8; violet areas). Previous work has shown that not all TE insertions are paramutated [10, 48, 56]. We therefore assumed a variable fraction of paramutable loci. It is unclear what determines whether or not an insertion may be paramutated but it is likely that the genomic neighborhood plays a role. For example, a TE insertion into a repetitive region may be more readily paramutated than an insertion in a gene. We thus assumed that the genomic site determines whether or not a locus is paramutable. As a consequence, all individuals in the population have the same set of paramutable loci. A ‘paramutable TE’ is a TE insertion in a paramutable locus which does not produce piRNAs, i.e. the TE has not (yet) been paramutated (fig. 1A: scenario 3; TE on the right). A ‘paramutated TE’ is a TE insertion in a paramutable locus that produces piRNAs, i.e. the TE has been paramutated (fig 1A: scenario 7; TE on the right). Since paramutations require both maternally deposited piRNAs and a TE insertion in a paramutable locus, a TE silenced in the mother may be reactivated in the offspring if the maternally deposited piRNAs but not the paramutable TE (nor a cluster insertion) is inherited (fig. 1A: scenario 5). This is in agreement with previous work which has shown that the maternally deposited component without chromosomal component is not sufficient to maintain the silencing of a reporter construct [19, 34]. As a consequence, we assume that a TE is active in all individuals not having at least one piRNA-producing locus, i.e. either an insertion into a piRNA cluster or a paramutated TE (fig. 1A: scenarios 1,3,5). One important consideration is that paramutations may maintain the silencing of an invading TE, but since they depend on maternally deposited piRNAs, they cannot explain the origin of the first piRNAs against an invading TE. Initially, we explored a scenario where a TE insertion in a piRNA cluster triggers the emergence of the first piRNAs against an invading TE (fig. 1A: scenarios 2,4). Later, we consider the case that siRNAs might activate the piRNA-based host defence [34].

**Figure 1:**
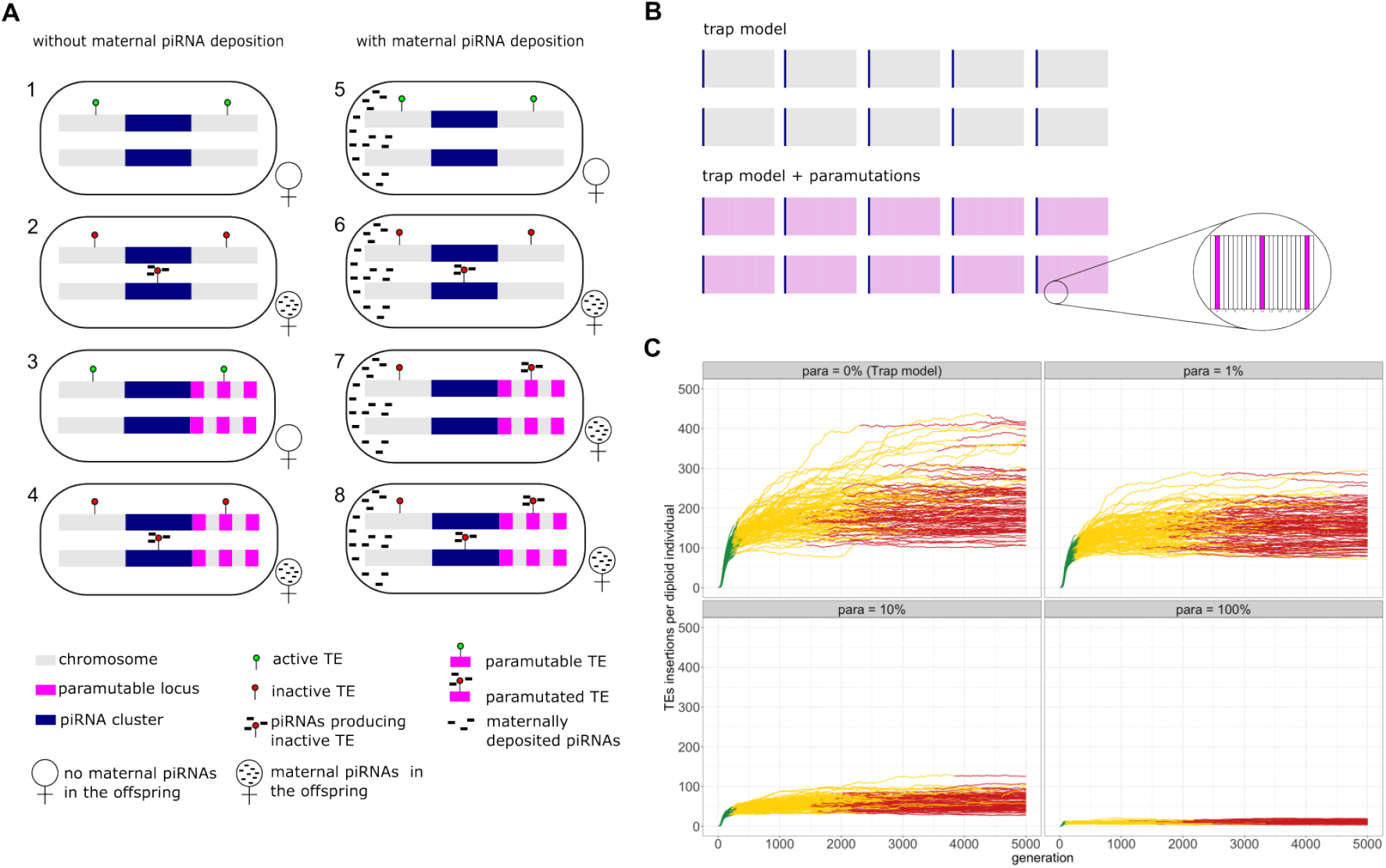
An overview of our simulations. A) Overview of model assumptions and the notation used in this work. B) Overview of the simulated genomes. We simulated five chromosomes, each with a piRNA cluster at the end. In the simulations with paramutations, the paramutable loci (pink) were evenly distributed in the genome. C) Abundance of TE insertions during TE invasions with different fractions of paramutable loci (para; top panel). Note that paramutations substantially reduce the number of TE insertions accumulating during a TE invasion. Colors indicate the three distinct phases of TE invasions (see text).

To explore the influence of paramutations on TE invasions, we performed individual-based forward simulations. With our novel simulator InvadeGO, developed in the Go programming language, we simulated TE invasions in panmictic populations of diploid organisms with discrete generations. Since paramutations depend on maternally transmitted piRNAs, we simulated two sexes (males and females) in contrast to our previous work where only hermaphrodites were simulated [24, 23]. The presence/absence of maternally deposited piRNAs was simulated as a binary trait. We thoroughly validated the novel simulator by comparing the observed results against theoretical expectations for all relevant population genetic forces, such as transposition, drift, recombination and selection (supplementary figs. S1-S4). Additionally, we validated the correct implementation of the genomic landscape, piRNA clusters and paramutations (supplementary figs. S5-S7). All simulations and analyses conducted in this work have been carefully documented with RMarkdown and deposited, together with the resulting figures, at GitHub (https://github.com/Almo96/Paramutations_TEs; see the md-files in the subfolders).

In this work, we simulated 5 chromosome arms with a length of 10Mb, a uniform recombination rate of 4cM/Mbp and piRNA clusters at the beginning of each chromosome (fig. 1B). We assumed that the paramutable loci are regularly distributed in the genome, e.g. with 10% paramutable loci a TE insertion at each 10th locus could be paramutated (fig. 1B).

Unless mentioned otherwise, we used the following default parameters: a transposition rate of *u* = 0.1, a population size of *N* = 1000, piRNA clusters with a size of 300kb (i.e. 3% of the genome) and 10% paramutable sites. Initially, we explored neutral TE insertions (*x* = 0.0) but later simulated more complex scenarios with negatively selected TEs. We triggered the TE invasions by distributing 100 insertions randomly in a population of 1000 individuals (population frequency *f* = 1/(2 · 1000)) to avoid the early stochastic stage of TE invasions, where TEs are frequently lost by genetic drift [29, 23].

We first asked whether TE invasions can be stopped under a trap model with paramutations. We define an invasion to be “stopped” (inactive phase) once a piRNA producing locus (either a cluster insertion or paramutated TE) becomes fixed in the population, since the TE can no longer be active in any single individual at any generation following fixation. Irrespective of the number of paramutable loci, 92 - 100% of the invasions were stopped after 5,000 generations and all after 10,000 generations (supplementary table S1; fig. 1C).

Next, we asked whether paramutations have an influence on the dynamics of TE invasions. To test this, we simulated TE invasions in populations with varying fractions of paramutable loci (0%, 1%, 10% and 100%; (fig. 1C top panel). Note that the scenario without paramutable loci (0%) corresponds to the classic trap model, which holds that TE invasions are solely controlled by TE insertions in piRNA clusters. Interestingly, we observed that paramutations exert a profound effect on TE invasions (fig. 1C), highlighting the importance of investigating the impact of paramutations on the dynamics of TE invasions.

### Effect of paramutations on TE invasions

To quantify the effect of paramutations on TE invasions, we relied on the three phases of TE invasions identified in a previous work [23]. In our previous work, we only considered insertions in piRNA clusters to define the three phases (no paramutations were considered). For this work, we extended the definition of the phases to incorporate paramutated loci. Thus, we define the three phases as follows; In the first phase, the rapid invasion phase, the TE may spread in the population largely uninhibited by the host defence (fig. 1C green). In the second phase, the shotgun phase, the spread of the TE is controlled by segregating piRNA producing loci (i.e. either a cluster insertions or paramutated loci). The onset of this phase is defined as the time when at least 99% of the individuals carry at least one piRNA producing locus (fig. 1C yellow). In the final phase, the inactive phase, the TE is inactivated by a fixed piRNA producing locus. The onset of the phase is the time when the first piRNA producing locus reaches fixation (fig. 1C red).

Based on these definitions, we found that paramutations significantly reduce the length of the rapid invasion phase (fig. 2A; Kruskal-Wallis rank sum test χ^2^ = 274.0, *p* = 2.2 · 10^-16^). Therefore, paramutations enable hosts to control TE invasions more rapidly. Paramutations only had a moderate effect on the length of the shotgun phase (fig. 2A; Kruskal-Wallis rank sum test χ^2^ = 16.7, *p* = 8.2 · 10^-4^). Next, we investigated the influence of paramutations on the abundance of TEs. Given that the time required to establish host control differs among replicate populations, we measured the TE abundance not at a fixed time point but rather at the onset of the shotgun and inactive phase. This corresponds to the moment when a TE invasion is initially brought under host control, either through segregating or fixed piRNA-producing TE insertions. We found that the abundance of the TEs dramatically decreased with increasing numbers of paramutable loci (fig. 2B; shotgun phase, Kruskal-Wallis rank sum test χ^2^ = 350.1, *p* = 2.2 · 10^-16^, inactive phase, Kruskal-Wallis rank sum test χ^2^ = 328.5, *p* = 2.2 · 10^-16^). The average number of piRNA cluster insertions per diploid individual also decreased with the fraction of paramutable loci (fig. 2C; shotgun phase Kruskal-Wallis rank sum test χ^2^ = 368.5, *p* = 2.2·10^-16^, Kruskal-Wallis rank sum test χ^2^ = 301.6, *p* = 2.2·10^-16^). Accordingly, the fraction of individuals with at least one cluster insertion also decreased with the fraction of paramutable loci (fig. 2D; shotgun phase Kruskal-Wallis rank sum test χ^2^ = 365.6, *p* = 2.2 · 10^-16^, inactive phase Kruskal-Wallis rank sum test χ^2^ = 281.0, *p* = 2.2 · 10^-16^). On the other hand, the number of paramutable TEs and the fraction of individuals with a paramutated TE increased with the abundance of paramutable loci (fig. 2E,F; number of paramutated TEs: shotgun phase, Kruskal-Wallis rank sum test χ^2^ = 380.0, *p* = 2.2 · 10^-16^, inactive phase, Kruskal-Wallis rank sum test χ^2^ = 280.97, *p* = 2.2 · 10^-16^, fraction of individuals with paramutated TE: shotgun phase Kruskal-Wallis rank sum test χ^2^ = 373.9, *p* = 2.2 · 10^-16^, inactive phase Kruskal-Wallis rank sum test χ^2^ = 311.9, *p* = 2.2 · 10^-16^). Our data show that paramutated TEs may, to some extent, compensate for cluster insertions, even though cluster insertions are necessary to initiate piRNA production. To more carefully investigate the effect of paramutations on the abundance of TEs, we performed simulations where we varied the fraction of paramutable loci in steps of 5% (fig. 2G,H); Both the abundance of TEs and cluster insertions declined rapidly with increasing numbers of paramutable loci until about 10-20% of the loci were paramutable (fig. 2G,H). Further increases in the number of paramutable loci only had a minor effect on the abundance of TEs, suggesting that around 10-20% paramutable loci are sufficient to markedly reduce the number of TEs accumulating during TE invasion.

**Figure 2:**
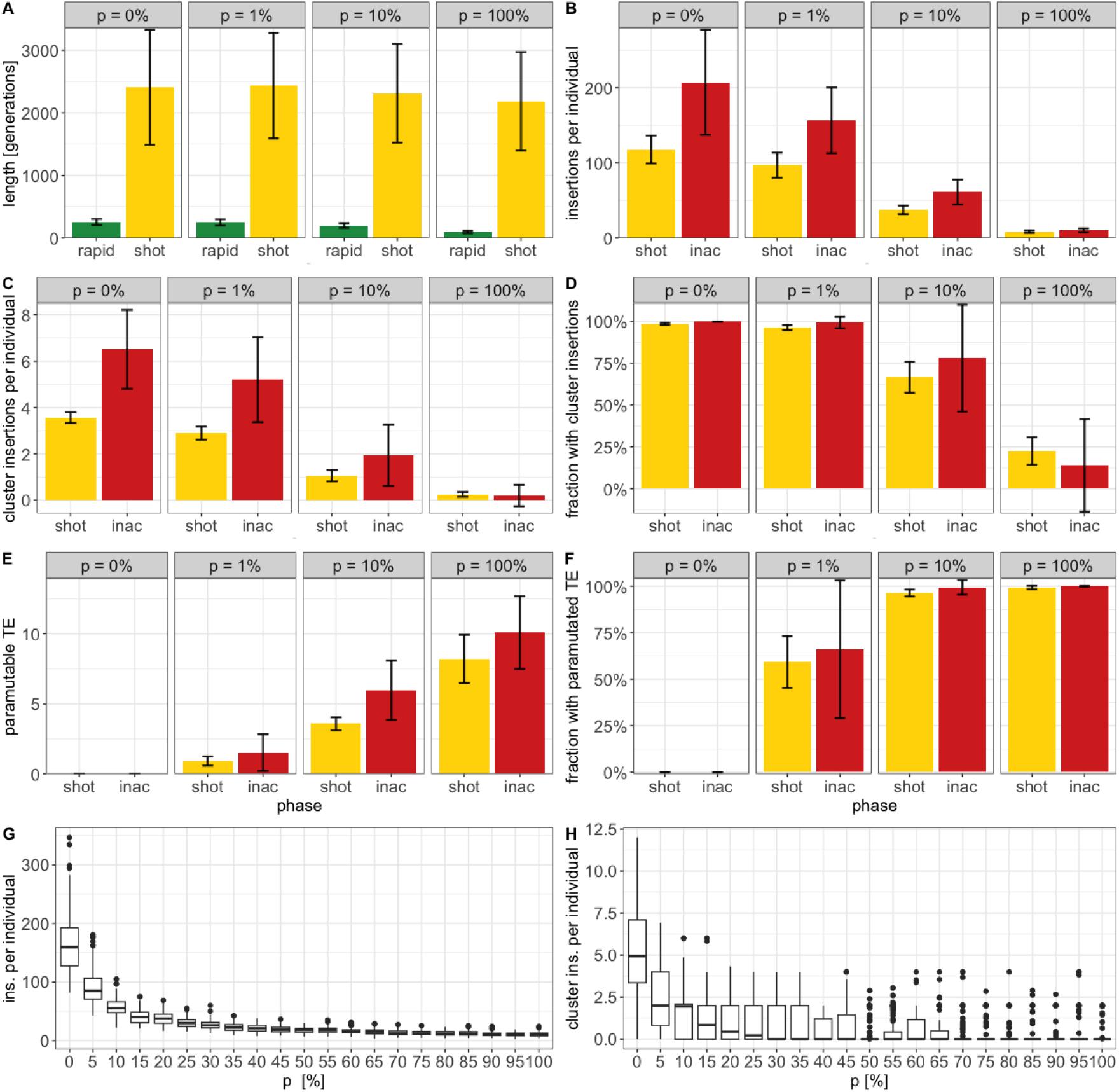
Effect of paramutations on different key properties of TE invasions. A) Length of the phase B) TE abundance per diploid individual at the start of the phase C) Cluster insertions per diploid individual at the start of the phase D) Fraction of individuals with at least one cluster insertion at the start of the phase E) Paramutable TEs per diploid individual at the start of the phase F) Fraction of individuals with at least one paramutated TE at the start of the phase. G) Effect of the fraction of paramutable loci (*p*) on the average number of TE insertions per diploid individual (at generation 5000). H) Effect of paramutable loci on the number of cluster insertions (at generation 5000).

Next, we explored the interaction between the size of piRNA clusters (in % of the genome) and the fraction of paramutable loci (supplementary fig. S8). In the absence of paramutable loci (i.e. trap model), increasing the size of piRNA clusters from 1% to 50% had little effect on the abundance of cluster insertions. In agreement with our previous work, about 4 cluster insertions were necessary to control TE invasion under the trap model, regardless of the cluster size (supplementary fig. S8A; [23]). However, when 10% paramutable loci were simulated, the size of piRNA clusters had a substantial effect: the number of cluster insertions required to silence an invasion increased from 0.3 with a cluster size of 1% to 4 when clusters account for 50% of the genome. In contrast, the number of paramutable TE insertions decreased from about 4 (cluster size 1%) to about 0.34 (cluster size 50%; (supplementary fig. S8B). Therefore, paramutations are most important when the size of piRNA clusters is small. This further supports our finding that paramutated TEs may, to some extent, compensate for cluster insertions.

In summary, we showed that paramutations speed up the silencing of TE invasions and reduce the amount of TEs accumulating during TE invasions. Additionally, paramutated TEs may compensate for cluster insertions. When paramutations are considered, the expected number of TE insertions in piRNA clusters is lower than under the classical trap model (without paramutations).

### Paramutations reduce the fitness burden of TEs

Motivated by our finding that paramutations reduce the number of TEs accumulating during an invasion, we next asked whether paramutations may also reduce the fitness burden generated by TE invasions. To explore this question, we introduced deleterious effects of TE insertions into the simulations (*x* > 0). To avoid an unlikely equilibrium state (TSC-balance [23]) where an invading TE is never silenced in all individuals in a population, we assumed that TE insertions in piRNA cluster are neutral (i.e. *x_cluster_* = 0). We simulated a linear fitness cost of TE insertions *w* = 1 — *x* · *n* where *w* is the fitness of an individual, *n* the number of TE insertions per diploid individual (outside of piRNA clusters) and *x* the negative effect of a TE. The invasion dynamics of TEs with deleterious effects differ substantially from the invasion of neutral TEs (supplementary fig. S9). For neutral TEs, copy numbers rapidly increase in early generations. In later generations, piRNA producing loci, which slow down the invasion, start to emerge, such that the TE abundance gradually reaches a stable plateau (supplementary fig. S9; [23]). For deleterious TEs, copy numbers also rapidly increase at early generations. At later generations, when the piRNA-based host defence emerges, the TE abundance declines as deleterious TEs are purged from populations by negative selection (supplementary fig. S9; [23, 21, 24]). This dynamic is reflected in the average fitness during the invasions of deleterious TEs. Initially, we observe a rapid fitness decline due to the accumulation of deleterious TE insertions (fig. 3A). At later generations, when the host defence is established, the average fitness rapidly recovers as deleterious insertions are purged from the population by negative selection. Therefore, TE invasions likely reduce host fitness solely temporarily (fig. 3A; [24, 21]). Here, we refer to the lowest fitness of individuals during a TE invasion as the ‘minimum fitness” (fig. 3A black arrow). The minimum fitness thus represents the maximal fitness burden generated by TE invasions. The minimum fitness may be a crucial factor determining the persistence of host populations as it has been argued that populations could go extinct if this temporary fitness reduction is too severe [24]. We found that paramutations have a marked impact on host fitness during TE invasions, where the fitness reduction during invasions is less severe in the presence of paramutable loci (fig. 3A). Paramutations significantly reduce the fitness cost of TE invasions (fitness cost = 1 - minimum fitness; fig. 3B).

**Figure 3:**
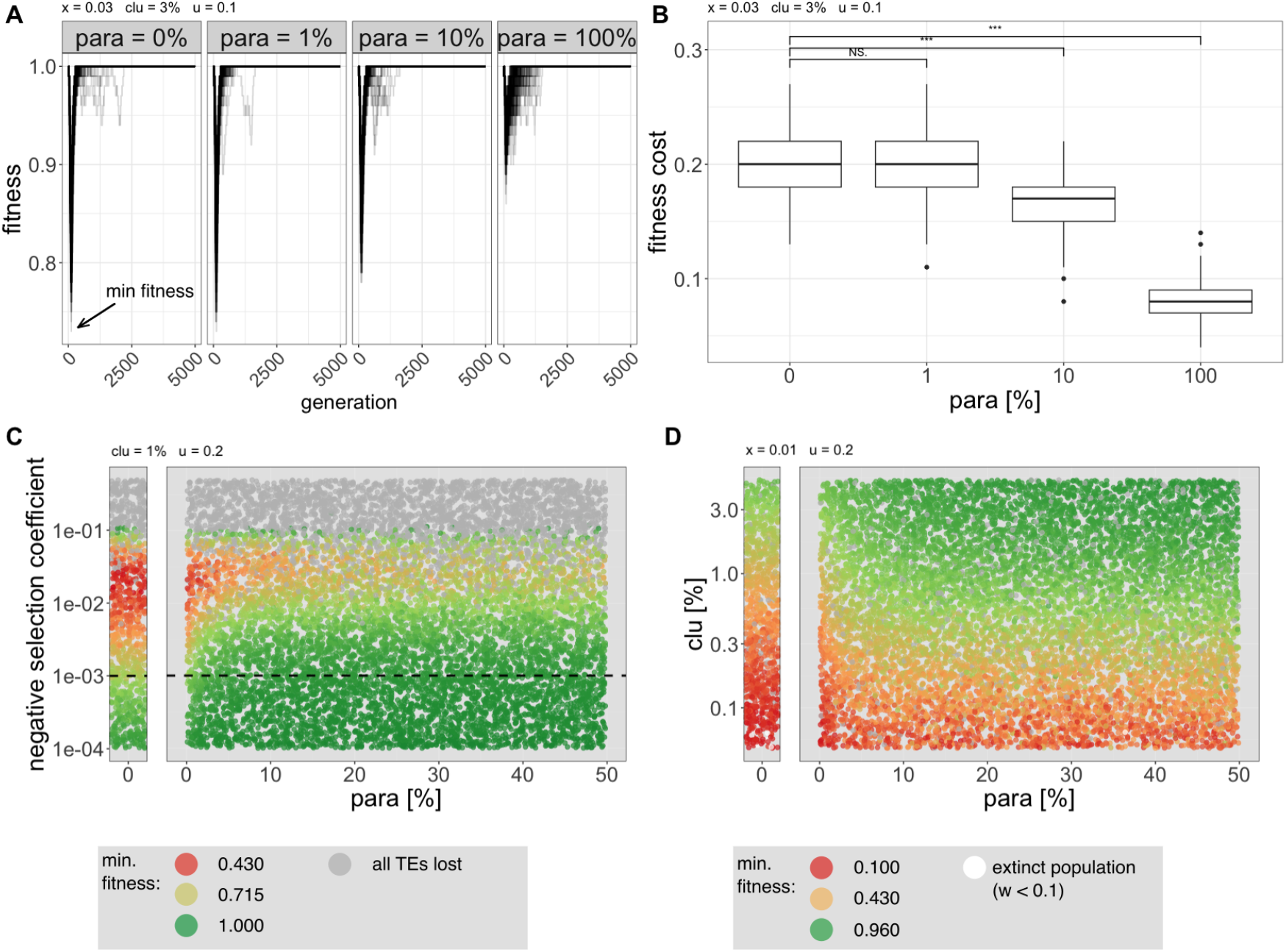
Paramutations reduce the fitness burden generated by TE invasions. A) Average fitness of individuals during TE invasions with different fractions of paramutable loci (top panel). The lowest fitness during the invasion (minimum fitness) is indicated. B) Paramutations reduce the fitness cost of TE invasions (fitness cost = 1 - minimum fitness). The significance of the fitness reduction was estimated with Wilcoxon rank sum tests (*** *p* < 0.001). C) Paramutations reduce the fitness cost of TE invasions, for many different negative effects of TEs. Each dot represents the result of a simulated TE invasion with a randomly chosen negative effect of TEs (y-axis) and a fraction of paramutable loci (x-axis). Colors indicate the minimum fitness during the invasion. Simulations which lost all TEs are shown in grey. Negative selection is not effective when *N* · *x* < 1 (dashed line). Additional data for simulations without paramutable loci (classic trap model) are shown in a separate left panel. D) Paramutations reduce the fitness cost of TE invasions for different sizes of piRNA clusters. Each dot represents the minimum fitness during a TE invasion where the size of the piRNA cluster (y-axis) and the fraction of paramutable loci (x-axis) were randomly chosen. Extinct populations (minimum fitness < 0.1) are shown in white.

The aforementioned simulations were performed with a deleterious TE effect of *x* = 0.03. However, the fitness effects of TEs remain an important open question [2] and the deleterious effect of TEs may vary among TE families, where for example long TE families may be more deleterious than short ones [44]. We thus aimed to systematically explore the impact of deleterious TE effects in combination with different fractions of paramutable loci on host fitness. To cover the parameter space, we performed 10, 000 simulations. For each simulation, we randomly picked a negative effect of TEs (between *x* = 0.0001 and *x* = 0.5) and a fraction of paramutable loci (between *para* = 0% and *para* = 50%). We performed additional simulations without any paramutable loci to more clearly show the expectations for the classic trap model (*para* = 0%; fig. 3C; left panel, trap model). After 5, 000 generations, we recorded the minimum fitness for each simulation. In agreement with previous work, we found that all TE insertions quickly got purged from populations when TEs are highly deleterious (here *x* > 0.1; 3C; [23, 24]). We found that over most of the explored parameter space, TE invasions led to a substantial fitness burden and that paramutations considerably reduced this fitness burden. The beneficial effect of paramutations was most pronounced when selection against TEs was effective (i.e. above the drift barrier where *N* · *x* > 1; fig. 3C; dashed line). Interestingly, about 10% paramutable loci are sufficient to markedly reduce the fitness burden generated by TE invasions (fig. 3C).

It is feasible that the beneficial effect of paramutations depends on the size of piRNA clusters. Therefore, we again used massive simulations to explore the effect of the size of piRNA clusters (0.1 — 4%) in combination with different fractions of paramutable loci (0 — 50%) on host fitness (fig. 3D). We performed additional simulations (1 , 000) to more clearly show the expectations under the classic trap model (0% paramutable loci; fig. 3D). We found that paramutations significantly reduced the fitness cost of TEs for all explored piRNA cluster sizes (fig. 3D; supplementary fig. S10). Again 10% of paramutable loci are sufficient to dramatically reduce the fitness burden of TEs.

In summary, we found that paramutations substantially reduce the fitness burden caused by TE invasions, where about 10% paramutable loci are sufficient to largely achieve the beneficial effect of paramutations.

### Epigenetically inherited piRNA production may be positively selected

Mediated by maternally deposited piRNAs, paramutations drive the conversion of TE insertions into piRNA producing loci. Once the first piRNAs against an invading TE have emerged, increasing numbers of TE insertions may be converted into paramutated TEs during TE invasions. We speculated that the piRNA-based silencing of TEs could spread in populations by a chain-reaction where paramutated TEs produce piRNAs that drive further paramutations in the next generation. Hence, we wondered whether paramutation dependent piRNA production (PDPP) could spread in populations in a non-Mendelian manner, similar to a gene drive. We aimed to construct a simple scenario that allows testing this hypothesis. We simulated a genome with five chromosomes (10Mbp) and a fixed TE insertion at a single paramutable locus (at the beginning of the first chromosome; fig. 4A). Consequently, each individual at any generation will carry a paramutable TE. The availability of paramutable TEs will, therefore, not be a limiting factor during the simulations. We used a transposition rate of *u* = 0.0, and a recombination rate of 4cM/Mbp but did not simulate piRNA clusters, Finally, irrespective of the sex, we initiated half of the individuals in the base population with maternally deposited piRNAs and the other half with no piRNAs (fig. 4A). If our hypothesis is correct and PDPP is spreading in a population similar to a gene drive, PDPP should become fixed in the majority of the simulated populations (if PDPP becomes fixed, all individuals in a population will produce piRNAs). On the other hand, if PDPP is solely subject to genetic drift, the fraction of populations where PDPP becomes fixed or lost should be similar. Our simulations show that the fraction of populations where PDPP was lost/fixed is roughly similar, demonstrating that PDPP is not *per se* spreading in populations (fig. 4B). This can be explained by the fact that only half of the individuals can propagate PDPP (females). Although males can inherit maternally deposited piRNAs, they cannot transmit them to the next generation. PDPP may only spread in populations like a gene drive if male gametes would also transmit piRNAs that could trigger paramutations in the next generation.

**Figure 4:**
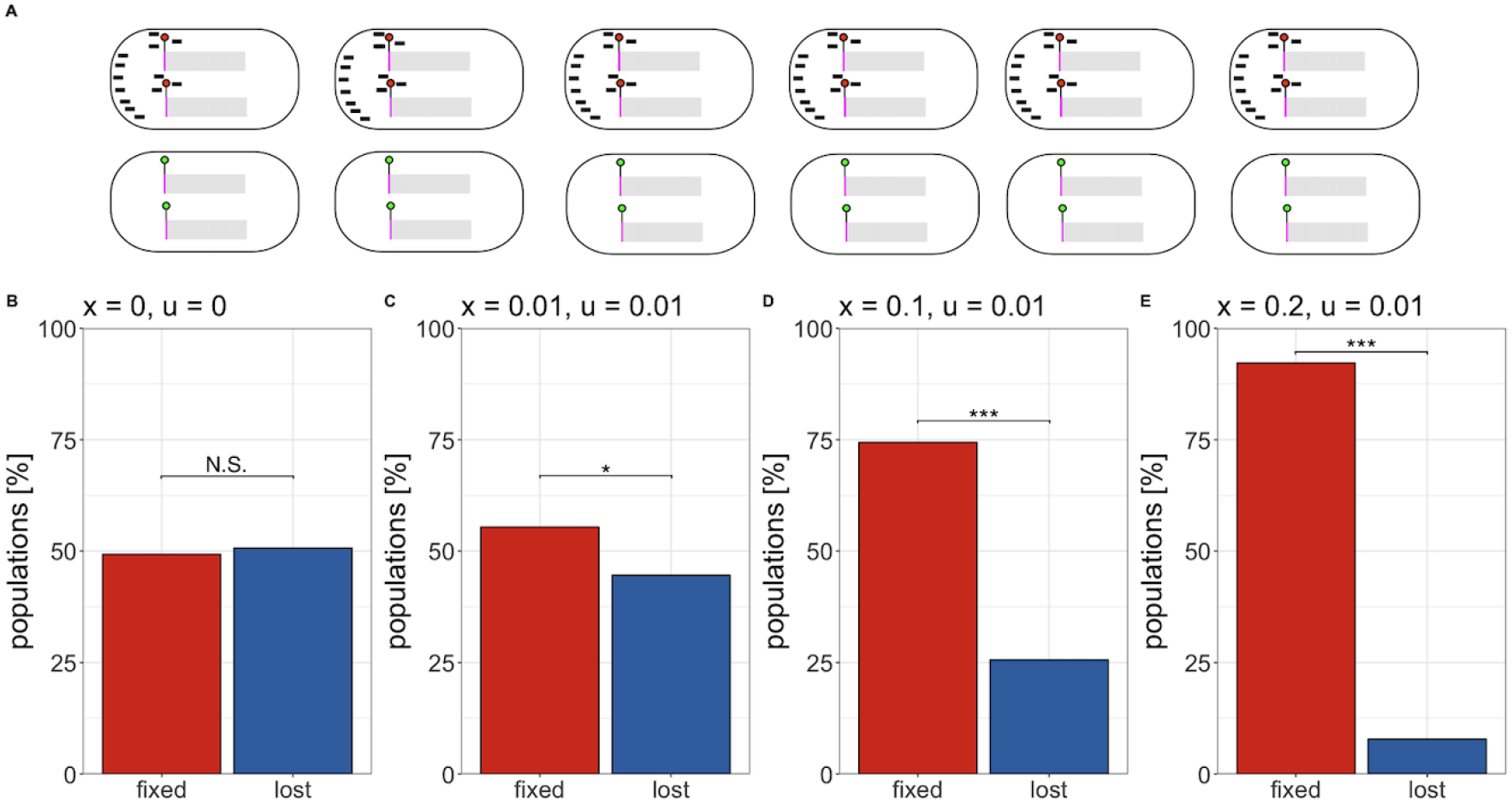
An epigenetically regulated trait - i.e. paramutation dependent piRNA production (PDPP) - may be positively selected. A) An overview of the simulated base population. We simulated a fixed TE insertion at a single paramutable locus (circle). Half of the individuals were assigned maternally inherited piRNAs, while the other half were not assigned any. B) Fractions of populations where PDPP is fixed or lost after 5000 generations. If PDPP is fixed, all individuals in a population will produce piRNAs. We simulated different negative effects of TEs (*x*) and different transposition rates (*u*). For neutral TEs (*x* = 0), we observed roughly equal ratios of populations where PDPP was fixed or lost, demonstrating that PDPP is not *per se* spreading in populations. However, when negative effects of TEs where introduced *x* > 0 the fraction of populations where PDPP was fixed increased, suggesting that the epigenetically inherited trait, PDPP, is positively selected. The significance was estimated with Chi-squared tests (* *p* < 0.05, *** *p* < 0.001).

Next, we wondered whether PDPP, an epigenetically inherited trait, could be positively selected. Individuals with an active host defence (e.g. PDPP) may accumulate fewer deleterious TE insertions than individuals without a host defence. These fitness differences may result in positive selection of PDPP. To test this idea, we simulated the same scenario as described before (fixed TE at paramutable locus; half of the individuals have piRNAs), with the sole difference being that we introduced deleterious effects of TEs (*x* > 0) and non-zero transposition rates (*u* > 0). In these scenarios, PDPP becomes fixed in most simulations, suggesting that PDPP is positively selected (fig. 4B).

In summary, we showed that an epigenetically inherited trait, paramutation dependent piRNA production (PDPP), may be positively selected as it reduces the fitness burden imposed by deleterious TE insertions.

### Paramutations may account for three important open problems with the trap model

Finally, we asked if paramutations could account for three important open problems with the trap model. First, several works showed that the number of TE insertions in piRNA clusters is lower than expected under the trap model. Studies that monitored P-element invasions in experimental populations, using Illumina short reads, found that P-element copy numbers stabilized after about 20-80 generations due to the emergence of piRNAs, but the number of P-element insertions in piRNA clusters at this stable plateau was lower than expected under the trap model [28, 27]. A study that monitored P-element copy numbers during invasions, with long-read data, also found that the number of cluster insertions were far lower than expected [47]. Furthermore, a recent study investigating the composition of piRNA clusters in different *D. melanogaster* strains, found that many TE families have fewer insertions in piRNA clusters than would be expected under the trap model [56]. Here, we demonstrated that the expected number of TE insertions in piRNA clusters decreases with increasing numbers of paramutable loci (fig. 2; supplementary fig. S8). Therefore, paramutated TEs could compensate for cluster insertions and paramutations may thus account for the insufficient number of cluster insertions observed in previous works [28, 27, 47, 56].

Second, a recent work found that deletion of three major piRNA clusters in *D. melanogaster* had no effect on the activity of the TEs [12]. The authors suggest that dispersed piRNA producing loci outside of piRNA clusters (i.e. likely paramutated TEs) may be responsible for producing the piRNAs that silence the TEs [12]. One vital question that remains unanswered in this work is how such dispersed piRNA producing loci arise in the first place. The generation of dispersed piRNA producing loci may require maternally deposited piRNAs and it is not yet clear how the first piRNAs complementary to an invading TE may emerge. In a commentary to this paper, Chen and Aravin [9] proposed that piRNA clusters could be crucial to trigger the initial host response but may become dispensable later on. Here, we explored this idea with a quantitative model. In particular, we test the hypothesis that paramutations may render piRNA clusters dispensable after a piRNA-based host defence emerged. Insertions in piRNA clusters may trigger the production of the very first piRNAs, that may drive the conversion of TE insertions into paramutated TEs. Once such paramutated TEs have emerged, piRNA clusters may be removed with little impact on the activity of the TEs.

To test this hypothesis, we generated a novel branch of InvadeGO (“remove-cluster”) that allows deleting a given number of piRNA clusters at a specified time (fig. 5 A). We again simulated populations of diploid organisms with 5 chromosomes and a piRNA cluster at the end of each chromosome (fig. 5 A). After 3000 generations, when the host defence has been largely established in all replicates, we removed between 0 and 4 out of 5 clusters from each individual in the population. We performed the simulations in a scenario with 0% (classic trap model) and 10% paramutable loci. Without paramutable loci, TE copy numbers dramatically increased in many replicates after the clusters were removed (fig. 5B; Wilcoxon rank sum test with average TE abundance after 5000 generations, clu –4 vs clu −0 *W* = 1388, *p* < 2.2 · 10^-16^). However, when 10% paramutable loci were introduced, no significant increase in TE copy numbers was observed even in the scenario where 4 out of the 5 clusters were removed (fig. 5B; Wilcoxon rank sum test with average TE abundance after 5000 generations, clu −4 vs clu −0 *W* = 4739, *p* = 0.52). Our simulations thus show that piRNA clusters could be important to trigger the host defence, but paramutated TEs render piRNA clusters dispensable once the piRNA-based host defence emerged.

**Figure 5:**
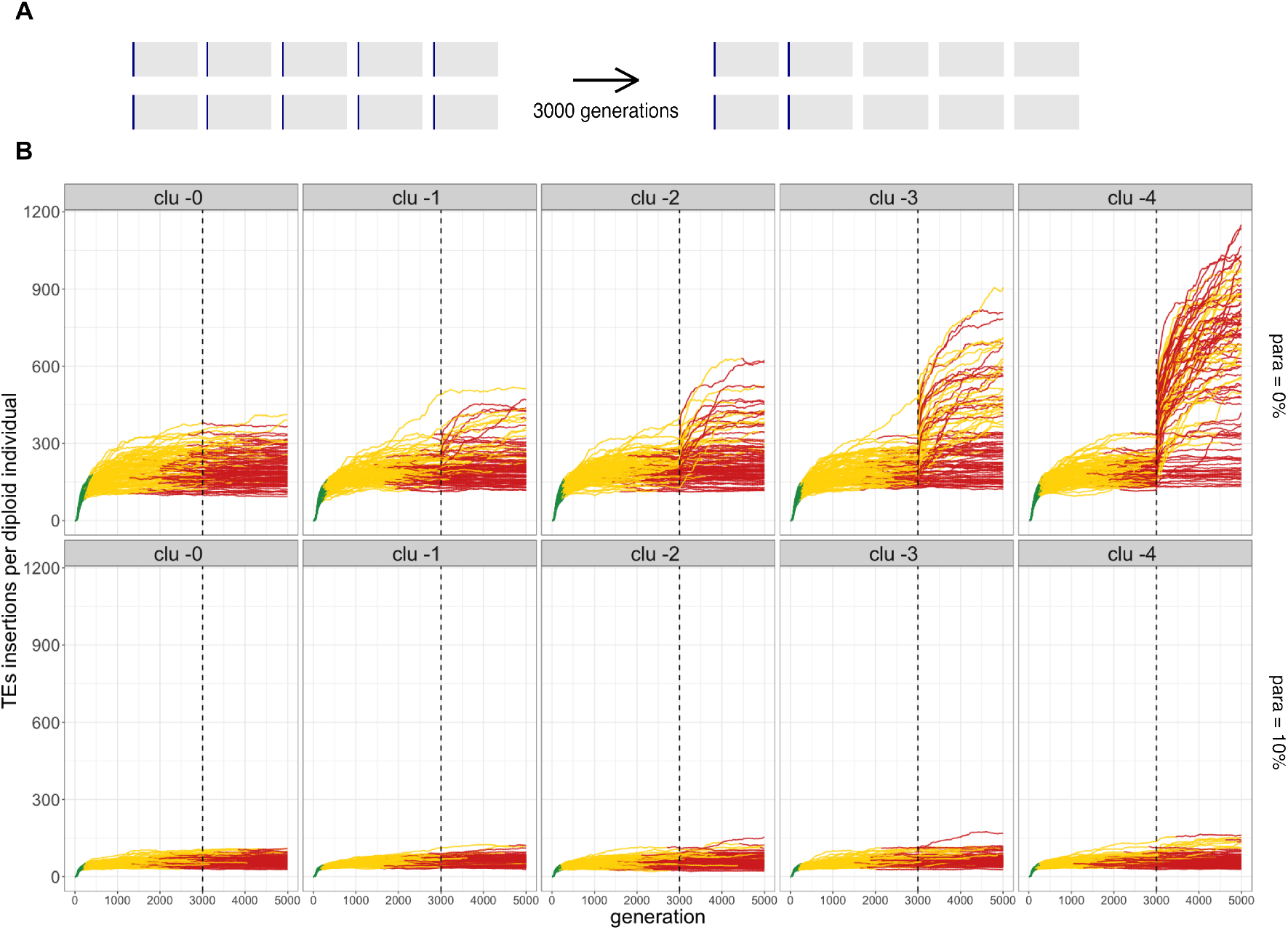
Paramutations render piRNA clusters dispensable once a piRNA-based host defence has been established. A) An overview of the simulated scenario. After 3000 generations, some piRNA clusters were removed from every individual in the population. B) Paramutations stabilize TE copy numbers when piRNA clusters are removed. Results are shown for different numbers of deleted clusters (top panel) and for simulations with 0% and 10% paramutable loci (right panel). Dashed lines indicate the time when the piRNA clusters were removed. Note that in the scenario with paramutations, the TE copy numbers remain largely stable after the clusters are removed. With paramutations, piRNA clusters are thus important to trigger the host defence but later become dispensable.

Finally, we tested whether paramutations can account for the discrepancy in the observed and the expected abundance of the different TE families in *D. melanogaster* [23]. Under the trap model, the size of piRNA clusters is a major factor determining the number of TE insertions accumulating during a TE invasion [23]. Apart from negative selection against TEs, all other investigated factors, such as transposition rate, genome size and recombination rate, only have a minor influence on the abundance of TEs [23]. If the size of piRNA clusters is known, quantitative predictions can be made about the expected abundance of TEs. In *D. melanogaster* where clusters account for about 3% of the genome, each TE family should have between 67.5 - 165 insertions per haploid genome (fig 6A). However, the actually observed abundance of most TE families is, with around 10 to 40 insertions per haploid genome, far lower than expected. It has been speculated that negative selection against TEs may account for this discrepancy [23]. Here, we explore an alternative hypothesis. We suggest that paramutations could also account for this discrepancy. To test this, we simulated TE invasions in populations of diploid organisms with piRNA clusters accounting for 3% of the genome, similar to dual-strand clusters in the germline of *D. melanogaster* [6]. Additionally, we either simulated 0% (classic trap model) or 10% paramutable loci. Finally, we compared the expected TE abundance (simulations) to the observed abundance of the different TE families in *D. melanogaster* (considering solely germline TEs; data from Kofler et al. [26]). As described previously, for the classical trap model, the observed TE abundance is much lower than expected (0% paramutable loci; expected: 67.5 - 165 insertions per haploid genome; 4 out of 98 TE families fit in this range; fig 6A; [23]). However, if 10% paramutable loci are used, the expected TE abundance fits much better with the observed abundance of the TE families (expected: 18.1 - 42.9 insertions per haploid genome; 29 out of 98 fit in this range; fig 6B). Therefore, a model with 10% paramutable loci and with piRNA clusters accounting for 3% of the genome, roughly captures the observed abundance of the different TE families in *D. melanogaster* (although some families with unusually high copy numbers can be observed; see discussion).

**Figure 6:**
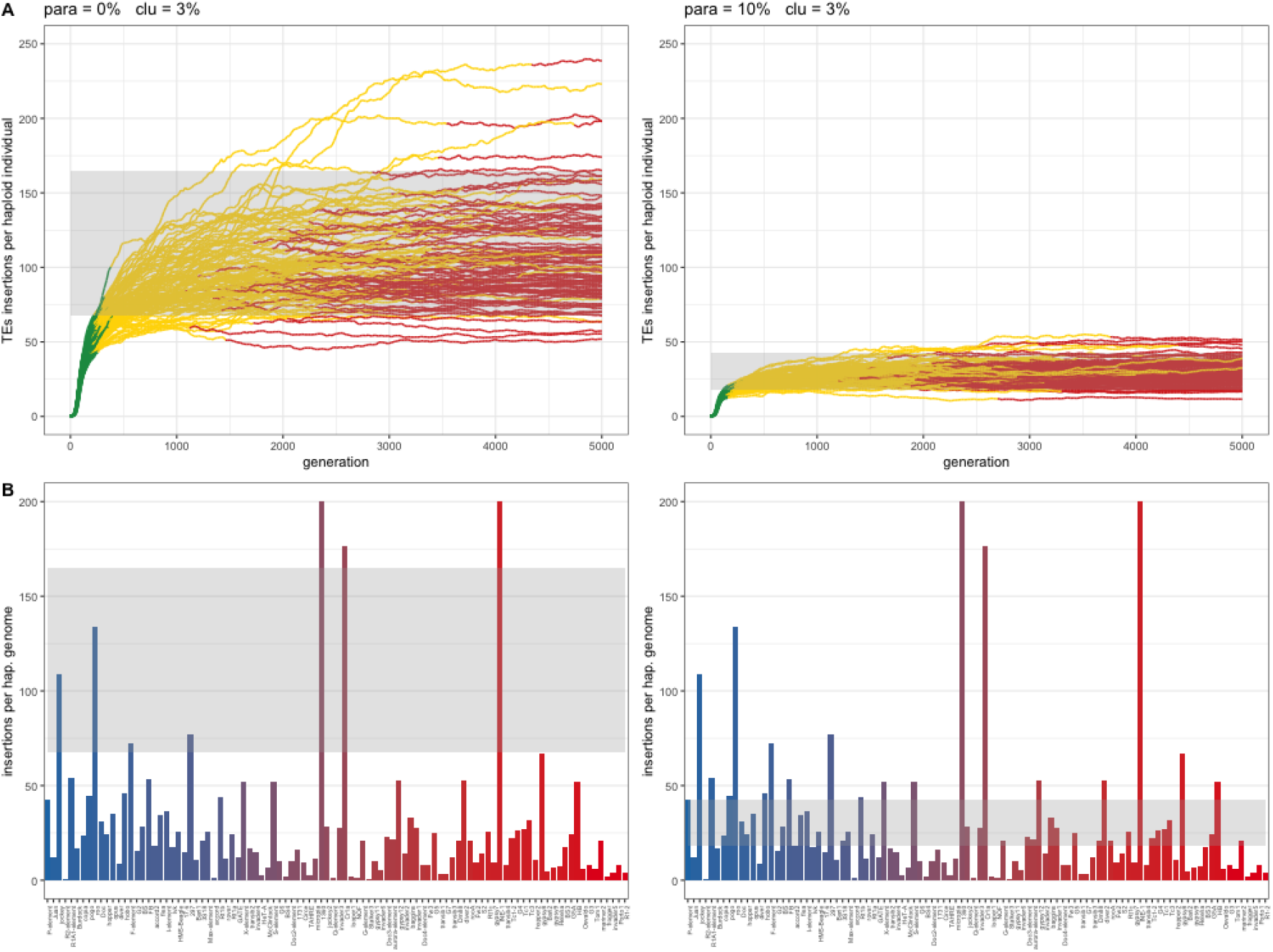
The trap model with paramutations predicts the observed abundance of TE families in *D. melanogaster* more accurately than a model without paramutations. A) Abundance of TEs during the simulated invasions with (10%) and without (0%) paramutable loci. In both scenarios, piRNA clusters account for 3% of the genome. B) Observed and expected abundance of TE families in *D. melanogaster*. Bar plots show the abundance of each TE family, where the colors of bars indicate the average population frequency (blue = 0.1, red = 1.0). The grey-shaded region highlights the expected abundance of the TE families based on the simulations. Note that the observed abundance of TEs is best captured by a model with paramutations. Data are from Kofler et al. [26].

In summary, we have shown that paramutations may account for three important open problems with the trap model. First, paramutations may compensate for cluster insertions, which could explain why the observed number of TE insertions in piRNA clusters is lower than expected under the classic trap model. Second, we showed that paramutations may render piRNA cluster dispensable once the host defence has been established, which could explain why deletion of three major clusters did not lead to an activation of TEs [12]. Third, paramutations will reduce the amount of TE insertions accumulating during invasions, which could explain why the abundance of many TE families in *D. melanogaster* is lower than expected under the classic trap model. A model with piRNA clusters (3% of the genome) and paramutations (10% of loci) largely predicts the abundance of TE families in *D. melanogaster*, even in the absence of negative selection against TEs.

### Impact of paramutations if siRNAs trigger the host defence

In agreement with the prevailing view, the trap model, we have assumed that a TE insertion into a piRNA cluster triggers the emergence of the first piRNAs required to initiate paramutations [4, 36, 60, 14, 59, 42]. However, it was recently suggested that siRNAs, generated from Dicer-2 mediated cleavage of dsRNA, may also initiate the emergence of piRNA producing loci [34]. Such dsRNA may be formed from sense and antisense transcripts of TEs [52, 34]. It is not obvious how to best capture this scenario with a simple model. Here, we opted to build the model based on the rate-limiting factor for dsRNA formation, the production of antisense RNA. Active TE families usually express ample sense RNAs such that sufficient amounts of the enzymes and other components necessary for transposition are generated. Antisense transcripts are usually more rare and are, therefore, most likely to be the bottleneck to any dsRNA formation. For example, we found that during an invasion of the P-element (a widely studied TE family) in experimental populations, about 7% of the P-element transcripts are antisense [47]. As proposed by Luo et al. [34], such antisense transcripts could result when a TE is inserted into a transcribed region (e.g. UTR or intron) in the opposite orientation to the gene. We model this notion by assuming that only TE insertions in some genomic regions can lead to the formation of antisense transcripts. We thus distributed siRNA-trigger-sites in the genome. We assumed that a TE insertion in a siRNA-trigger-site will lead to the production of antisense mRNA, that leads to the generation of siRNAs, which in turn drives the conversion of a TE insertion at a paramutable locus into a piRNA producing locus. Since TE insertions into many genes could likely trigger the production of antisense mRNA, we assumed that trigger-sites are more evenly distributed in the genome than piRNA clusters. In contrast to insertions in piRNA clusters, we assumed that insertions in siRNA-trigger-sites cannot themselves produce piRNAs (nor be converted into piRNA producing loci). This is in agreement with Luo et al. [34], who showed that a siRNA-producing hairpin, which activates piRNA production in trans, is not by itself producing piRNAs. The first piRNAs complementary to an invading TE will therefore emerge in individuals having i) at least one insertion in a siRNA-trigger-site and ii) at least one insertion in a paramutable locus.

Under our siRNA-model, between 90 and 100% of the invasions were stopped after 5,000 generations, and all invasions were stopped after 10,000 generations (supplementary table S2). The dynamics of TE invasions under this model are similar to the trap model. After an initial increase, TE copy numbers reach a stable plateau in all replicates (supplementary fig. S11). Again similarly to the trap model, three phases of TE invasions can be distinguished. First, TE copy numbers rapidly increase in the population (rapid invasion phase), next the invasion is silenced by segregating paramutated TEs (shotgun phase), and finally, the invasion is silenced by a fixed paramutated TE (supplementary fig. S11). Interestingly, once the proportion of trigger sites exceeds a certain minimum (≥ 1%), the amount of trigger sites has little impact on the abundance of TEs accumulating during an invasion (supplementary fig. S11). Paramutations also reduce the fitness burden generated by TE invasions under this model (supplementary fig. S12). Finally, we found that the observed abundance of different TE families in *D. melanogaster* can also be predicted under the siRNA model (assuming 10% paramutable loci and 3-30% trigger sites; S13).

To summarize, we found that the siRNA model, which assumes that insertions at some sites may trigger paramutations, is a viable alternative to the trap model. All TE invasions were stopped even in the absence of piRNA clusters and negative selection against TEs. piRNA clusters are, therefore, dispensable. Finally, paramutations again reduce the fitness burden of TEs, and the abundance of TE families in *D. melanogaster* can also be roughly predicted by a siRNA model.

## Discussion

Based on computer simulations with our novel tool InvadeGO, we show that paramutations are an important factor governing the dynamics of TE invasions. Paramutations limit the number of TE insertions accumulating during invasions, reduce the fitness burden of TEs, render piRNA clusters dispensable, and may account for the discrepancy in the observed and the expected TE abundance in *D. melanogaster*. Moreover, paramutation dependent piRNA production (PDPP), an epigenetically inherited trait, may be positively selected.

Our conclusions are based on several assumptions: i) transposons are spreading in populations at a given rate (*u*) ii) insertion sites of TEs are randomly distributed in the genome iii) piRNAs silence TEs iv) maternally deposited piRNAs induce paramutations v) some, but not all, TE insertions outside of piRNA clusters may be paramutated vi) an insertion into a piRNA cluster (or a siRNA-trigger-locus) may initiate the production of the first piRNAs complementary to an invading TE. All the aforementioned assumptions are well supported by the literature. First, the multiplication of TEs in genomes and populations has been reported by many studies[41, 5, 54, 28, 22, 1]. Second, although some TEs, such as the P-element, have an insertion bias [50, 25], little insertion bias could be found for most of the investigated TE families in *D. melanogaster* [18]. Third, there is little doubt that piRNAs repress the activity of TEs [6, 15, 30, 49, 42]. Fourth, our assumption that maternally deposited piRNAs may induce paramutations is also well supported by multiple works [10, 31, 16, 8]. Fifth, several previous studies have shown that some, but not all, of the TE insertions outside of piRNA clusters may be converted into piRNA producing loci [48, 56]. It is an important open question which genomic, epigenetic, and cellular factors determine whether or not a TE insertions may be paramutated. The TE family is likely one important factor since the fraction of paramutated TEs varies among the families [56]. The genomic context is likely also important. For example, Shpiz et al. [48] suggested that paramutations may happen next to transcribed genes, and de Vanssay et al. [10] suggested that a tandem arrangement of several TE insertions may be necessary for paramutation. It is conceivable that the composition of maternally deposited piRNAs and even the environment are also important factors influencing whether a locus may be paramutated (e.g. temperature [8]). Possibly the most important question is which events trigger the emergence of the very first piRNAs complementary to an invading TE. Here, we assumed that either an insertion in a piRNA cluster or siRNA-induced paramutations may initiate piRNA production. Several prior studies have shown that insertion of an artificial sequence into a piRNA cluster will lead to the generation of piRNAs complementary to this artificial sequence [39, 20, 34]. For example, Luo et al. [34] found that insertion of a GFP-reporter in a major piRNA cluster of *D. melanogaster*, 42AB, leads to the generation of piRNAs complementary to GFP. It is, therefore, reasonable to assume that an insertion into piRNA cluster can trigger the *de novo* formation of piRNAs. It was recently suggested that siRNAs could also initiate the formation of novel piRNA-producing loci and thus trigger the production of piRNAs [34]. Such siRNAs may, for example, result from cleavage of dsRNA consisting of sense and antisense transcripts of TEs. We modelled this mechanism by assuming that the expression of antisense transcripts will be the bottleneck in the formation of dsRNA and that only TE insertions at specific genomic sites will yield antisense transcripts (i.e. siRNA-trigger-sites). For example, antisense transcripts of TEs may result from TE insertions in transcribed regions in the opposite direction to the gene [34, 52]. However, there are still many unknowns about this mechanism: Are insertions in transcribed regions the main source of antisense transcripts of TEs? Which cellular processes are involved in the siRNA-mediated initiation of piRNA production? Thus, it is not clear how well our siRNA model captures the process of siRNA-mediated initiation of piRNA production. In any case, our simulations show that the siRNA-model as implemented in this work is a viable alternative to the trap model, as TE copy numbers stabilize after the initial rapid invasion in our siRNA-model.

We show that PDPP, an epigenetically inherited trait, may be positively selected. Individuals that produce piRNAs will accumulate fewer deleterious TE insertions than individuals without piRNAs, and therefore PDPP may be beneficial. This raises the question as to whether PDPP leaves footprints in populations that could be detected. As PDPP is an epigenetically inherited trait, it will not necessarily leave footprints in the genome [40]. This is illustrated by our simulations where selection on PDPP was observed, although the only TE insertion that could produce piRNAs was fixed in the population (fig. 4). It is, however, conceivable that some genomic loci, like the cluster insertion that triggered PDPP, could hitchhike with PDPP. Such an indirect increase in the frequency of some alleles may be detectable at the genomic level. Directly determining whether PDPP is positively selected is currently challenging because it requires estimating whether piRNA production is spreading more rapidly in populations than would be expected under a null mode (neutral expectations). Apart from the technical difficulty of measuring piRNA production in many individuals during TE invasions, inferring the expectations under the null model is also challenging, especially given that it is not yet entirely clear which events trigger the emergence of the piRNAs complementary to an invading TE.

We demonstrated that a model with paramutations captures the abundance of most TE families in *D. melanogaster* more accurately than a model without paramutations (fig. 6). Nevertheless, we find that some TE families, like roo and jockey, have much higher copy numbers than expected under the model with paramutations. It is unclear why some TE families accumulate such extremely high copy numbers. One possible solution comes from recent works which monitored P-element (a DNA transposon) invasions in experimental populations. Usually, the P-element is rapidly and consistently silenced in all experimental populations by the emergence of a piRNA-based host defence [27, 47]. However, Selvaraju et al. [47] found that in one replicate population, the host-defence could not be established, even after many generations, which also led to the accumulation of unusually high P-element copy numbers. Since this only happened in a single replicate, the failure to establish the host defence is likely a stochastic event. A delay or a failure in the establishment of host control over an invading TE could therefore be a possible explanation for the highly abundant TE families in *D. melanogaster*. An alternative explanation for the families with unusually high copy numbers could be an insertion bias of the TEs [50]. For example, an insertion bias into promoter regions may result in a low probability of jumping into piRNA clusters. TEs with such a bias could then accumulate large numbers of insertions before a sufficient number of cluster insertions appeared.

Paramutations limit the number of TE insertions accumulating during an invasion and thus reduce the fitness burden of TEs. Paramutations may therefore be beneficial to organisms. In agreement with this, we found that PDPP may be positively selected. This raises the question as to whether paramutations evolved as a distinct mechanism to robustly control TEs. An alternative explanation could be that paramutations are just a byproduct of the mechanisms responsible for maintaining the location of piRNA clusters across generations. The position of piRNA clusters in the next generation is, at least partially, determined by maternally deposited piRNAs [31]. Mediated by maternally deposited piRNAs, silencing chromatin marks necessary for the formation of piRNA clusters are installed [31]. While these piRNAs will be important to maintain the location of piRNA clusters across generations, they may also, as a side-effect, induce paramutations at dispersed TE insertions outside of piRNA clusters.

It has been shown that only some, e.g. around 5 to 10%, of the TE insertions, may be paramutated into piRNA-producing loci [56, 10, 38]. Why are not more or fewer insertions paramutated? It is feasible that paramutations are costly to organisms. Paramutations of TEs may, for example, reduce the expression level of neighbouring genes [48, 37]. Hence, paramutations may interfere with gene expression, and selection may thus act to reduce the frequency of paramutations. On the other hand, too few paramutations could also be deleterious to hosts. We showed that paramutations reduce the burden of TEs accumulating during invasions. Since individuals with many paramutations will accumulate fewer TEs, selection may act to increase the frequency of paramutations. However, if paramutations are primarily responsible for maintaining the location of piRNA clusters across generations (see above), then selection may act to increase the frequency of paramutations indirectly. If the frequency of paramutations decreases, the stable inheritance of piRNA clusters across generations may be jeopardized, potentially leading to the reactivation of certain TEs. As this TE activity may be deleterious, selection may act to maintain stable inheritance of piRNA clusters and thus, indirectly, also maintain high frequencies of paramutated loci. Overall, it is conceivable that the frequency of paramutations is in a balance between two opposing evolutionary forces acting to decrease and increase the number of paramutated TEs. It is interesting to note that in *D. melanogaster*, the fraction of paramutated TEs is approximately 5-10%, which coincides with the range where our simulations suggest that the marginal benefit of paramutations is the highest (i.e. a higher frequency of paramutated TEs would only have a minor effect on the fitness burden caused by TEs; fig. 3C,D; [56]).

In summary, we showed that paramutations have a substantial impact on the dynamics of TE invasions and may thus be a crucial component of the host defence against selfish DNA.

## Materials and Methods

### Simulation software

To simulate TE invasions with paramutations, we implemented a novel simulation software in the Go programming language: InvadeGO (v0.2.3). Several algorithms implemented in InvadeGO are inspired by our previous simulation tool, Invade, which is implemented in Java [23]. InvadeGO performs individual-based forward simulations of TE invasions in populations of diploid organism using discrete (non-overlapping) generations. A TE insertion is modelled as position (integer) in the half-open interval [0, *g*), where *g* is the genome size. The TE insertions in a haploid genome are thus modelled as a list of integers (i.e. the genomic insertion sites). A diploid individual carries two separate lists of insertion sites. Each chromosome occupies a unique non-overlapping territory in the genomic interval [0, *g*), such that each TE insertion is part of exactly one chromosome. The piRNA clusters occupy sub-regions of the chromosomes, such that a TE insertion may be part of either none or one piRNA cluster. Recurrent sites, such as paramutable loci and the siRNA-trigger-sites are modelled using the modulo operation. For example, with *x*%10 == 3 the sites 3, 13, 23… would be paramutable loci (except for sites in piRNA clusters). Maternally deposited piRNAs are modelled as a binary trait, where each individual may have (*m* > 0) or not-have (*m* = 0) maternal piRNAs. Each individual has a fitness *w*, which solely depends on the number of TE insertions *w* = 1 — *xn*, where *x* is the negative effect of a single TE insertion and *n* is the number of TE insertions per diploid individual. The fitness determines the mating probability (i.e. fecundity selection). We simulated two sexes, where “males” may only mate with ‘females”. Each parent generates a single gamete that is passed to the offspring. To create a gamete, first recombination and random assortment among chromosomes are simulated and then novel transposition events are introduced into the recombined gamete. We assumed that TEs multiply with a given transposition rate *u*, which is the probability that a TE insertion will generate a novel insertion in the next generation. A transposition rate of zero (*u* = 0) was used for individuals carrying a piRNA producing locus (either a cluster insertion or a paramutated TE). To avoid excessive computation times, we calculated, for each gamete, the number of novel insertion sites based on a Poisson distributed random variable with λ = *u* * *n*/2. Novel insertion sites were randomly chosen in the genomic interval [0, *g*). If a site was already occupied, the novel insertion was ignored. The sex was randomly assigned to each offspring. Only females transmitted the maternal piRNA status to the offspring.

InvadeGO allows providing a wide range of different parameters, such as the number of chromosomes, the size of the chromosomes, the size of the piRNA clusters, the distribution of the paramutable loci and of the siRNA-trigger-sites, the recombination rate, the transposition rate, the population size, the number of generations, the number of TE insertions in the base population, the negative effect of TEs and a flag indicating whether or not cluster insertions are selectively neutral. For the base population, it is also feasible to provide a file that contains for each individual the positions of the TE insertions, sex, and the maternal piRNA status.

To test whether piRNA clusters are dispensable after the piRNA-based host defence was established, we developed a novel branch of InvadeGO (remove cluster; v.br.rc-0.2.3) that allows removing a given number of piRNA clusters at the specified time.

InvadeGO was thoroughly tested with unit-tests (all tests can be run with the command *go test* ./…). Furthermore, we validated the correct implementation of all relevant population genetic forces, such as drift, recombination, selection and transposition (supplementary figs. S1 - S7).

### Simulations and data analysis

To minimize the parameter space, we performed simulations with default conditions and varied the parameter of interest. For our default conditions, we used a genome consisting of 5 chromosomes each having a size of 10Mbp (--genome MB:10,10,10,10,10), a recombination rate of 4cM/Mbp (--rr 4,4,4,4,4), 10% paramutable loci (--paramutation 10:1), piRNA clusters with a size of 300kbp at the beginning of each chromosome (-cluster kb:300,300,300,300,300; i.e. 3% of the genome), a transposition rate of 0.1 (--u 0.1), population size of 1000 (--N 1000) and a base population with 100 randomly distributed TE insertions (--basepop 100). Initially, we simulated neutral TE insertions (--x 0) but later explored more complex scenarios with negatively selected TEs. To cover the parameter space in two dimensions (fig. 3C,D), we used Python scripts that launched about 10, 000 simulations with randomly chosen parameter combinations. All data analysis was done in R [45] and data were visualized with ggplot2 [55].

## Author contributions

RK and AS conceived the work. RK designed and implemented InvadeGO. AS performed simulations and analyzed the data. RK and AS wrote the paper.

## Acknowledgments

This work was supported by an Austrian Science Fund (FWF) grant (P35093) to RK. ChatGPT, a language model developed by OpenAI, and Google Translate contributed to polishing the english of some difficult sections. We thank Filip Wierzbicki and Matthew Beaumont for comments and all members of the Institute of Population Genetics for feedback and support.

## Data availability

InvadeGO is available under the GNU general public license, version 3 at GitHub (https://github.com/RobertKofler/invadego). The simulations and the data analysis were documented with RMarkdown [58] and have been made publicly available, together with the resulting figures, at GitHub https://github.com/Almo96/Paramutations_TEs.

