## Supplement for "The impact of paramutations on the invasion dynamics of transposable elements"

December 2022

### Supplementary figures

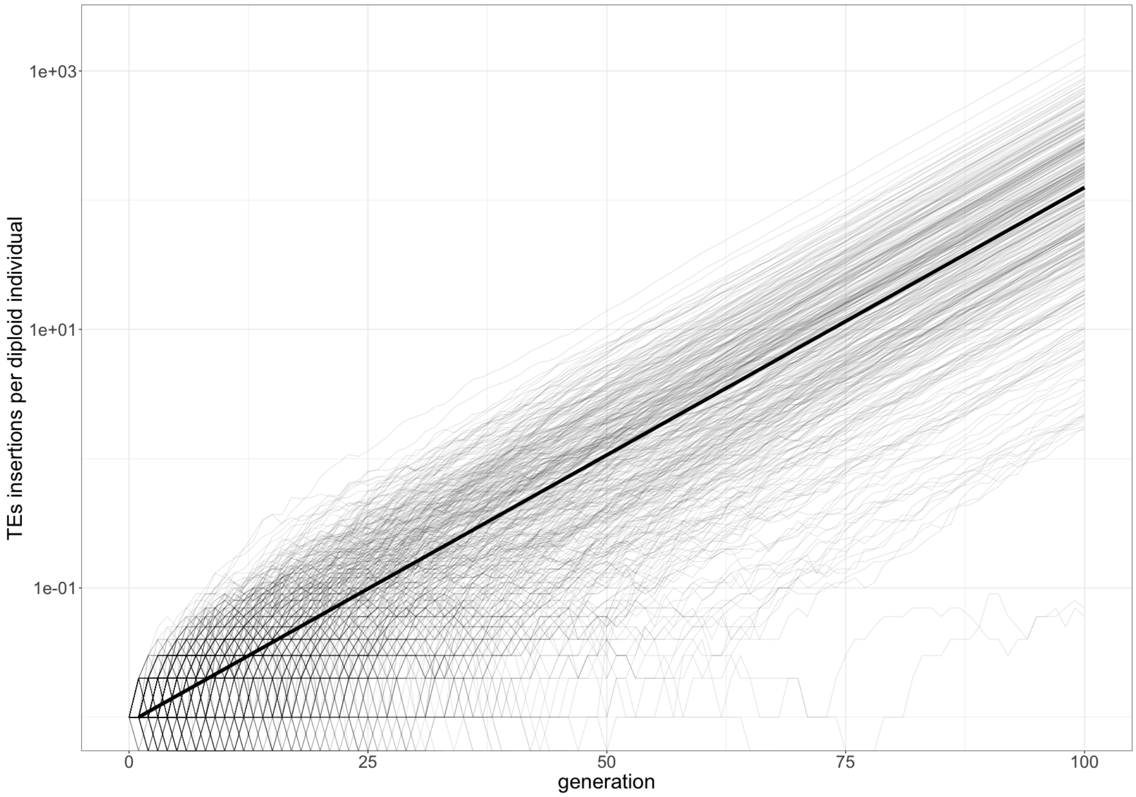


Figure 1: Validation that transposition is accurately simulated with InvadeGO. We tested whether the expected exponential growth of TE copy numbers in populations without host defence is accurately reproduced. We simulated 500 TE invasions with a transposition rate of $u = 0.1$ in populations of a size $N = 1000$. We started the invasion with 10 randomly distributed TE copies in the populations. Note that the observed TE copy numbers ($c$) follows closely the expectations ($c_t = c_0 \cdot (1 + u)^t$; bold line).

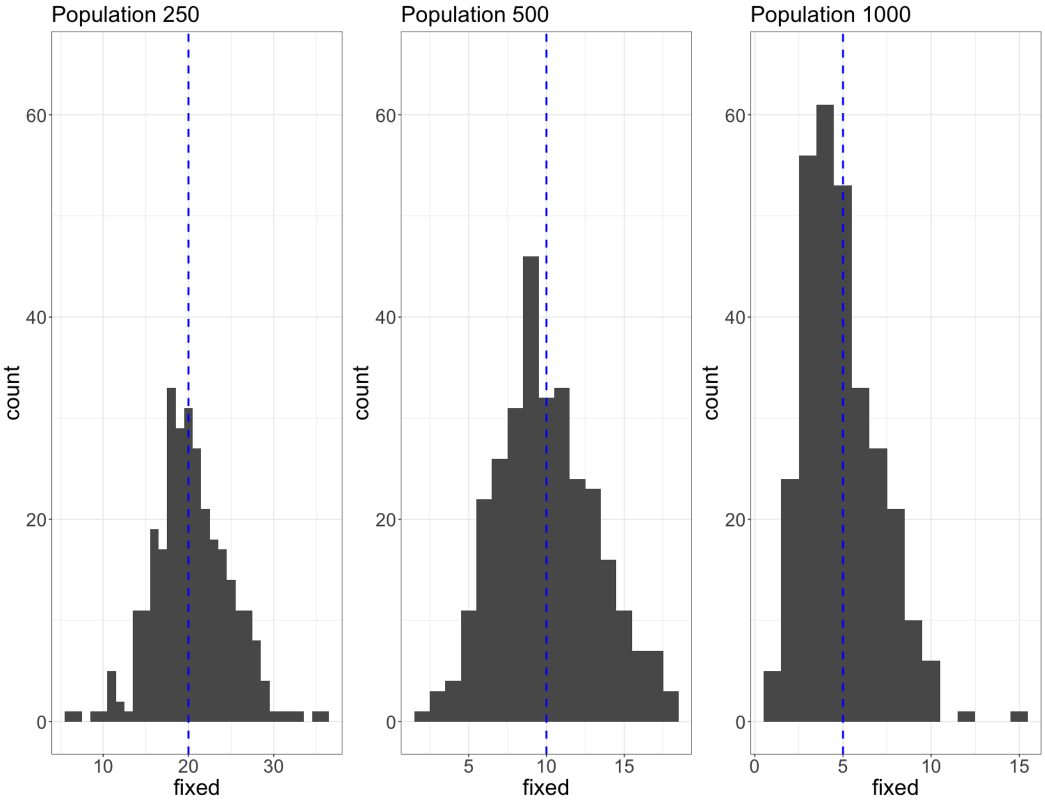


Figure 2: Validation that genetic drift is accurately simulated with InvadeGO. We relied on the basic population genetics insight that the probability of fixation of a neutral singleton is $p = 1/2N$. When $n$ loci are segregating as singletons the expected number of loci fixed by drift is $e = np$. To test this we distributed $10,000$ neutral TE insertions at random positions in populations with different population sizes ($N = 250$, $N = 500$, $N = 1000$). Importantly we used a transposition rate of $u = 0$ to prevent the emergence of additional TE copies. For each population size we used 300 replicates and populations were simulated for $20,000$ generations. In our scenario the expected number of fixed TE insertions is $e = 10000/2N$ (the dashed blue line). Note that the observed number of fixed TE insertions fits well with the expectations.

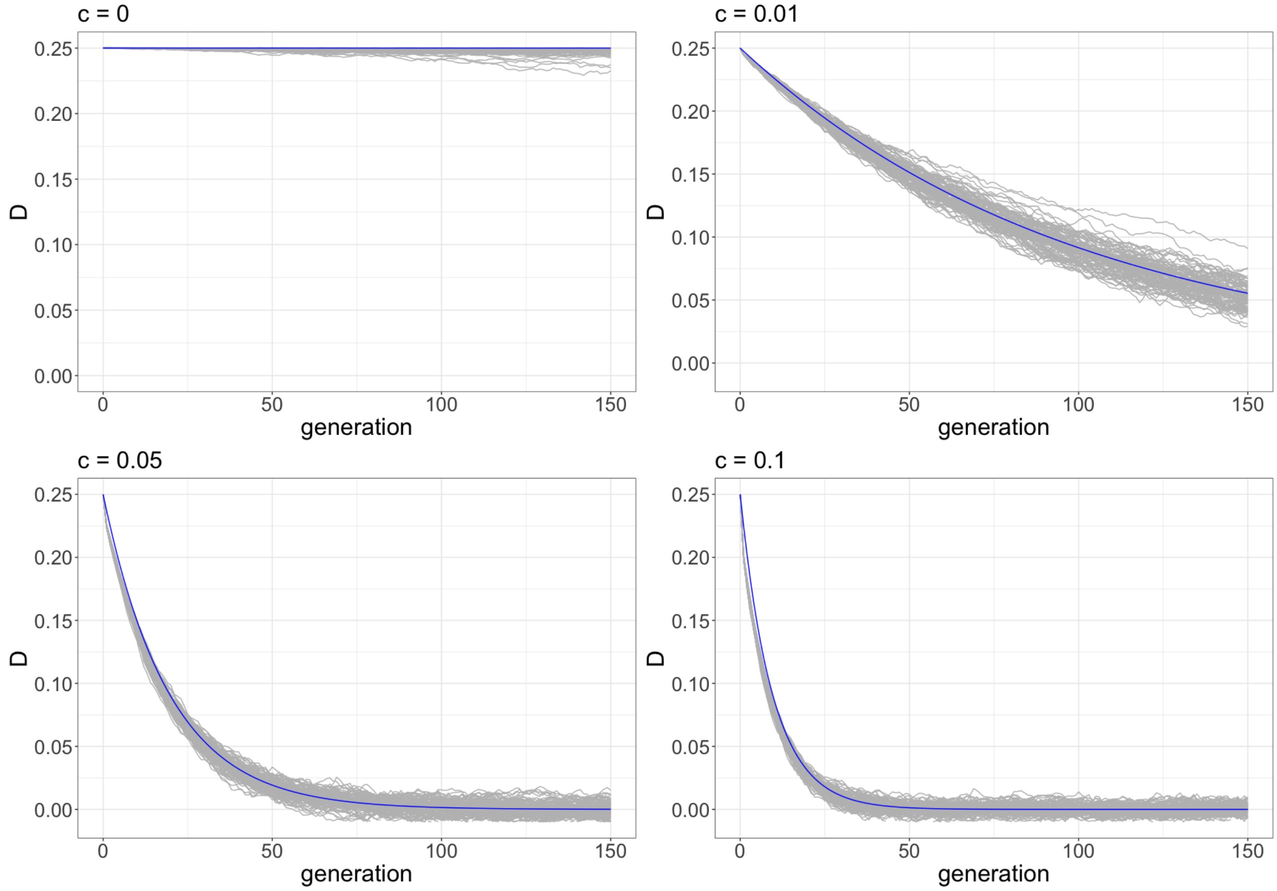


Figure 3: Validation that recombination is accurately simulated with InvadeGO. We tested whether the observed decay of linkage disequilibrium ($D$; grey lines) matches the theoretical expectations $D_t = D_0(1-c)^t$ (blue line) where $c$ is the recombination rate. For the base populations we simulated two neutral TE insertions with a population frequency of $f = 0.5$ in perfect linkage ($D = p_{AB} - p_A p_B = 0.5 - 0.5 \cdot 0.5 = 0.25$). Furthermore we used a population size of $N = 10000$ and a transposition rate of $u = 0$. We simulated four different recombination rates and used 100 replicates for each. The LD decay observed in the simulations agrees well with theoretical expectations.

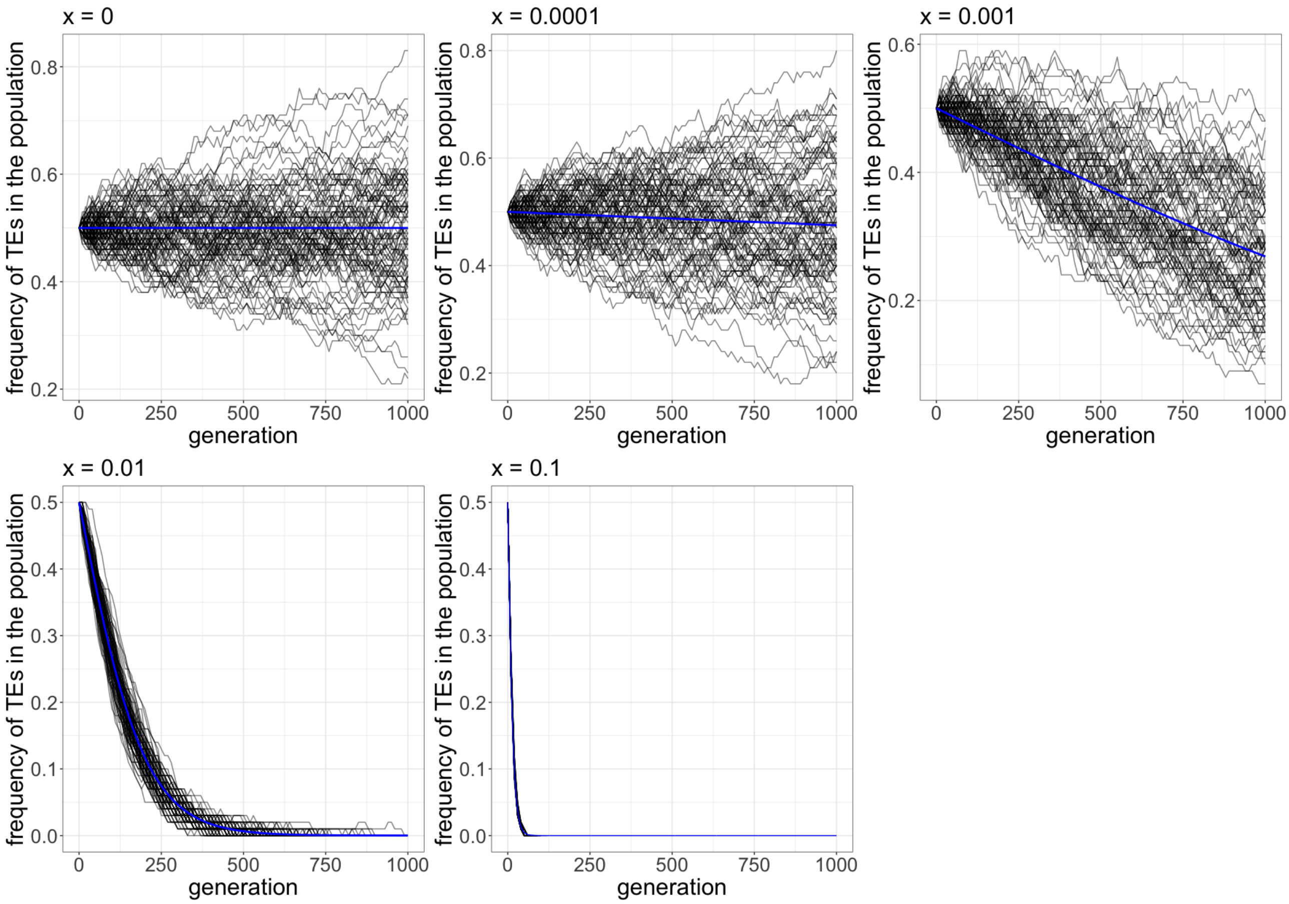


Figure 4: Validation that negative selection is accurately simulated with InvadeGO. We tested whether the observed trajectories of selected TEs matches the theoretical expectations. For a negatively selected TE with a population frequency of $p_t$ and a negative effect $x$ the allele frequency at the next generation $p_{t+1}$ can be computed as $p_{t+1} = p_t^2 w_{AA} + p_t(1-p_t)w_{Aa}/\overline{w}$ where $\overline{w}$ is the average fitness and $w_{AA} = 1-2x$ and $w_{Aa} = 1-x$ is the fitness of the homozygous and heterozygous TE insertion, respectively. We simulated a TE insertion with an allele frequency of $p_0 = 0.5$ (in Hardy-Weinberg equilibrium) in a population with size $N = 1000$. We simulated five different negative effects of TEs ($x$) and used 100 replicates for each. The simulations with $x = 0$ represent the neutral scenario. According to population genetic theory an allele should behave similarly to the neutral scenario when $N_e \cdot x < 1$ (i.e. $x = 0.0001$). Note that the trajectories of the negatively selected TE insertions agree with theoretical expectations.

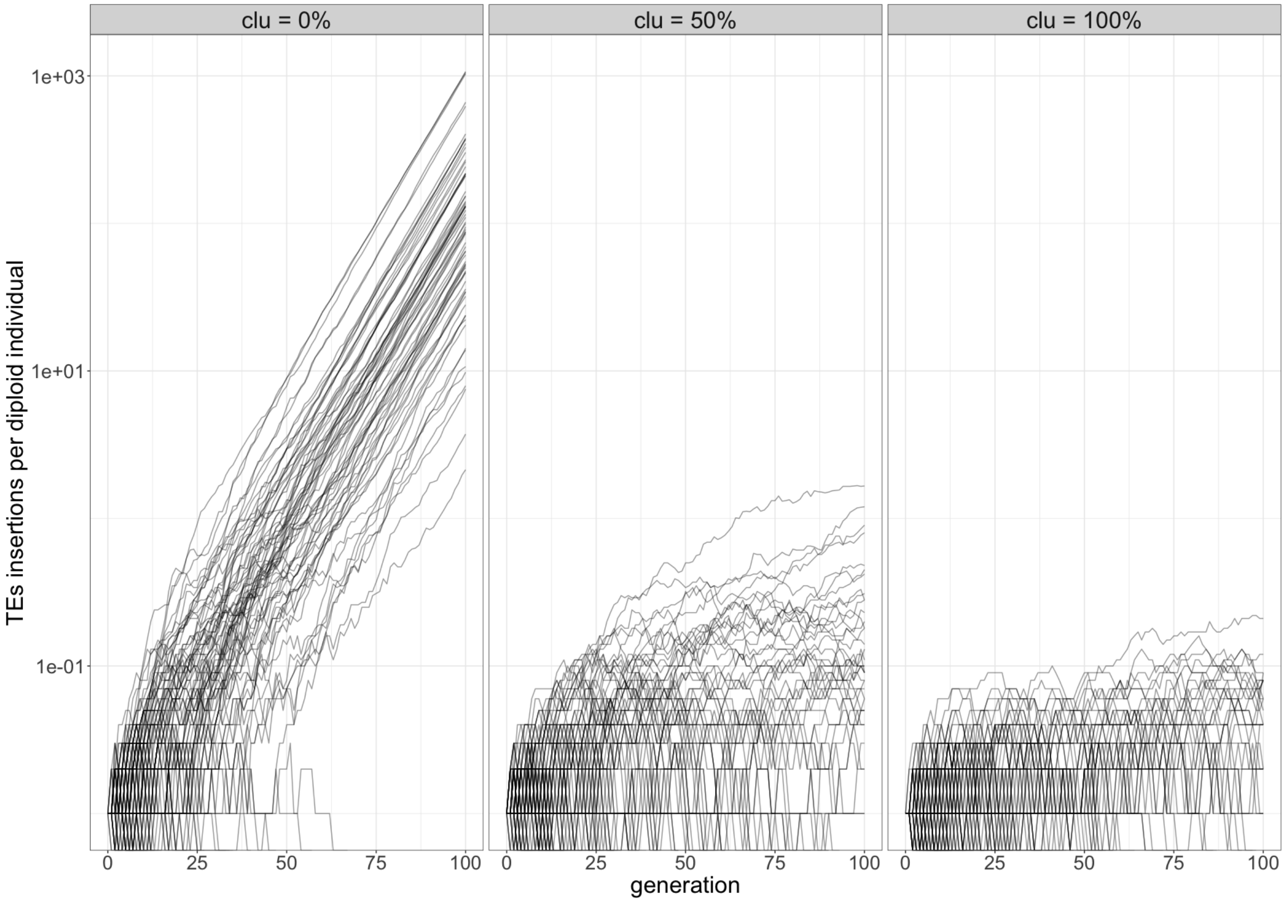


Figure 5: Validation that the effect of piRNA clusters is accurately simulated with InvadeGO. We simulated a single chromosome of 1Mb and a piRNA cluster at the end of the chromosome. piRNAs clusters either account for 0%, 50% and 100% of the chromosome. For each scenario we simulated 100 TE invasions with a transposition rate of $u = 0.1$ in populations of size $N = 1000$. We started the TE invasions with 10 randomly distributed TE copies in the populations. As expected TE copy numbers increased exponentially in the simulations without a piRNA clusters. Note that piRNA clusters markedly slowed down the copy number increase of the TE.

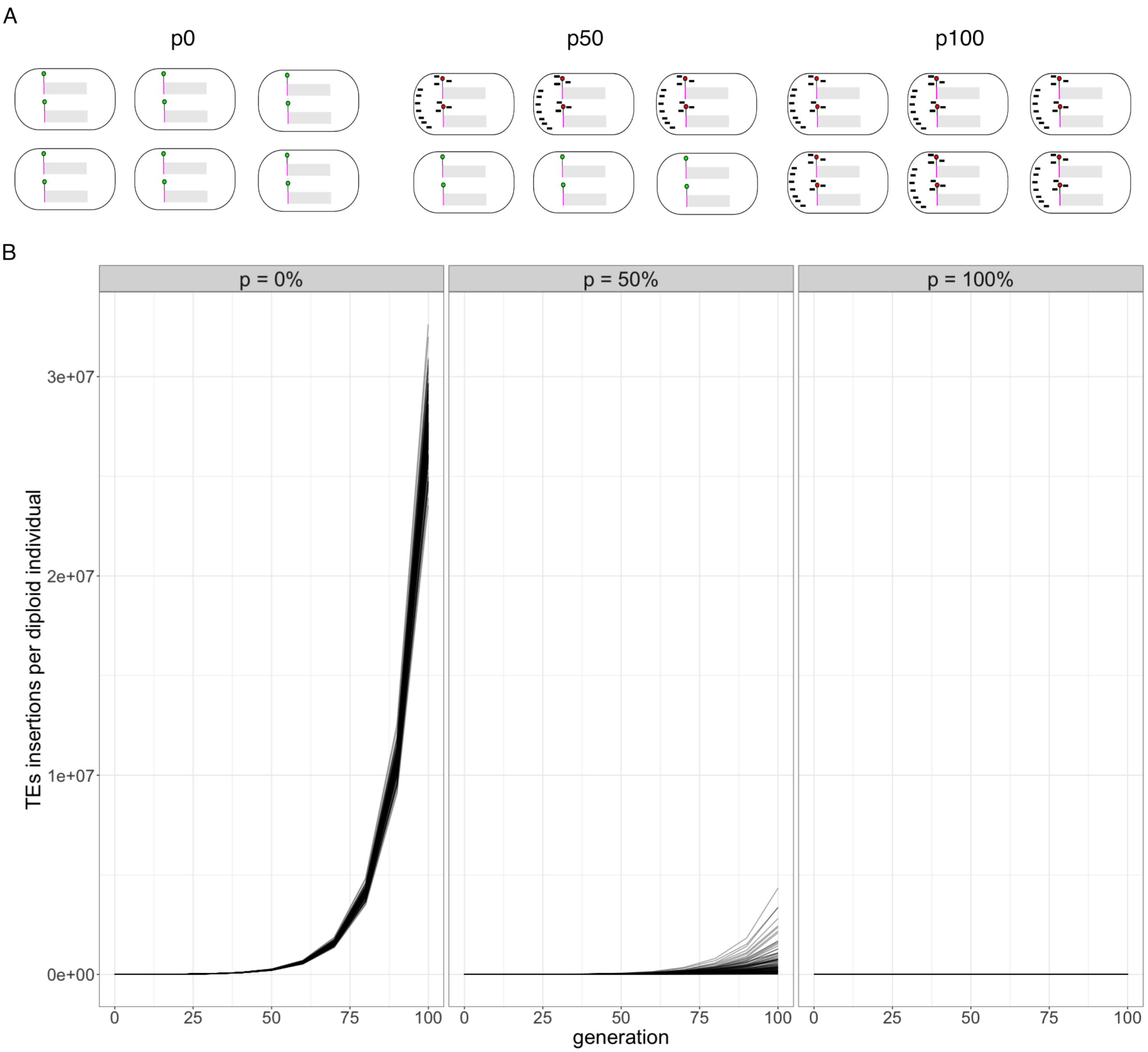


Figure 6: Validation that paramutations are correctly simulated with InvadeGO. A) Overview of the simulations. We simulated 1000 individuals, 1 chromosome of length 1Mb without piRNA clusters, 100 replicates and a fixed TE at a single paramutable locus (position 1). The transposition rate was $u = 0.1$. Different fractions of the individuals in the base populations were initiated with maternal piRNAs ( 0%, 50% and 100%). B) Average number of TE insertion per diploid individual during TE invasions, dependent on the fraction of individuals with maternal piRNAs in the base population (top panel). Note that maternal piRNAs reduce the amount of TEs accumulating during the invasions.

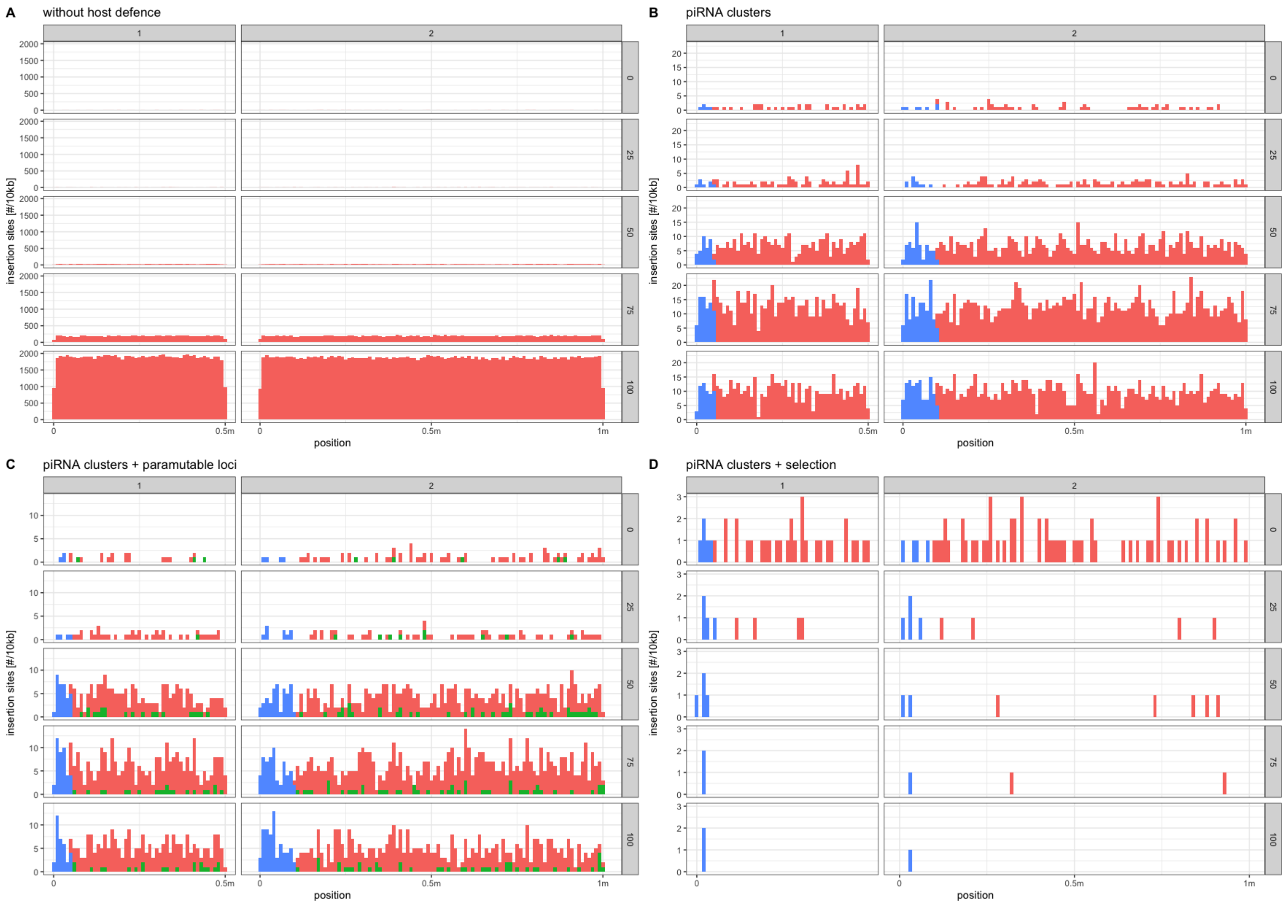


Figure 7: Validation that the genomic landscape and the random distribution of novel insertion sites is accurately simulated with InvadeGO. Simulations were performed for a diploid organism having two chromosomes (top panel) with an unequal size (500kb and 1000kb). The population size was $N = 1000$ and the transposition rate was $u = 0.1$. All plots show the number of insertion sites in 10kb windows at different generations during the TE invasion (right panel). Colors indicate the different genomic regions blue: piRNA cluster, green: paramutable loci, red: rest of the genome A) Simulation without host defence. Note that insertion sites are equally distributed in the two chromosomes. B) Simulation with piRNA clusters (100kb) at the 5' end of each chromosomes. Note that the cluster insertions are located at the end of the two chromosomes. C) Simulations with piRNA clusters and paramutable loci (at each 10th position outside of piRNA clusters). Note that paramutable loci are solely found outside of piRNA clusters and that paramutable loci are equally distributed over the chromosomes. D) Simulation with piRNA clusters and negative selection. We assumed that solely TE insertions outside of piRNA clusters are negatively selected. Note that all TE insertions, except for the insertions in piRNA clusters, are gradually lost from the population.

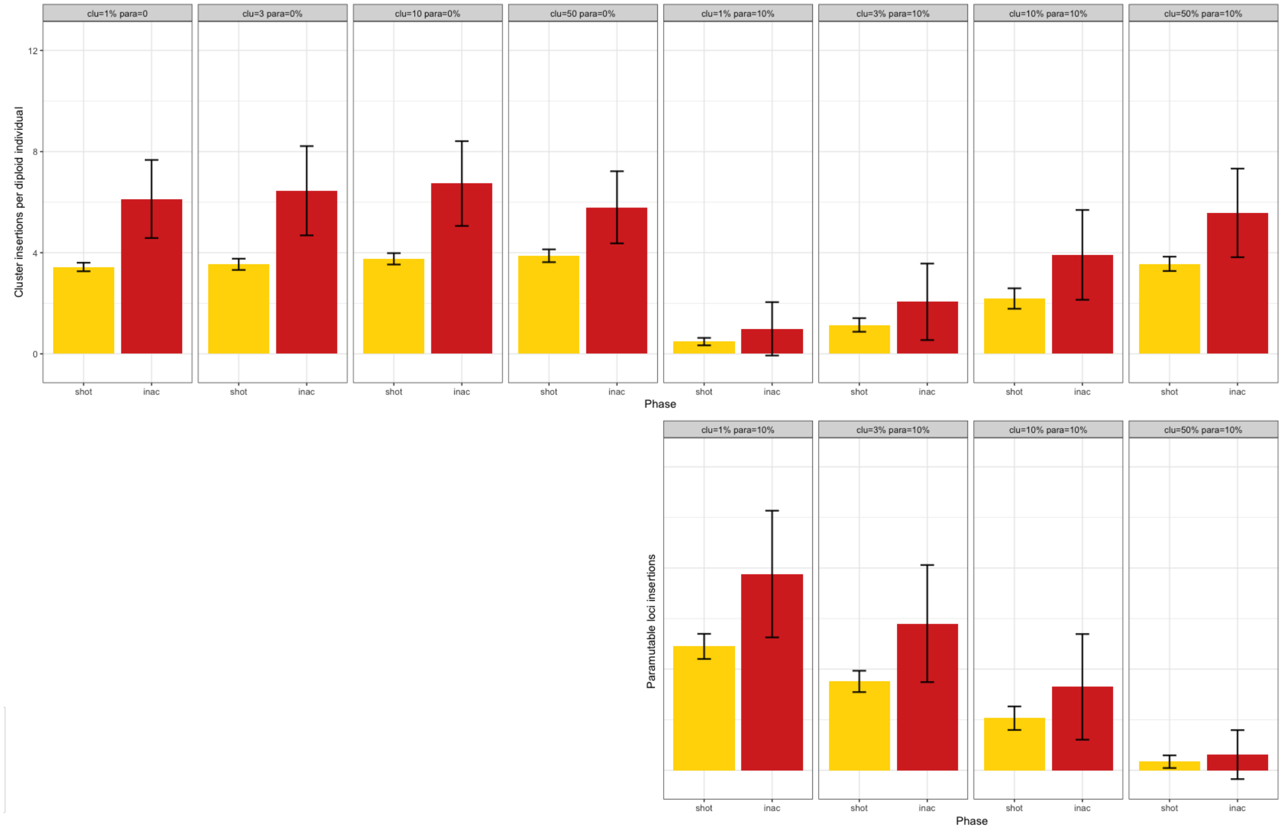


Figure 8: Interaction between the size of piRNA clusters ($clu$) and fraction of paramutatable loci ($para$). A) Number of cluster insertions. B) Number of paramutated TEs. Note that the number of paramutated TEs increases with decreasing size of piRNA clusters, which suggests that paramutated TEs may, to some extent, compensate for cluster insertions.

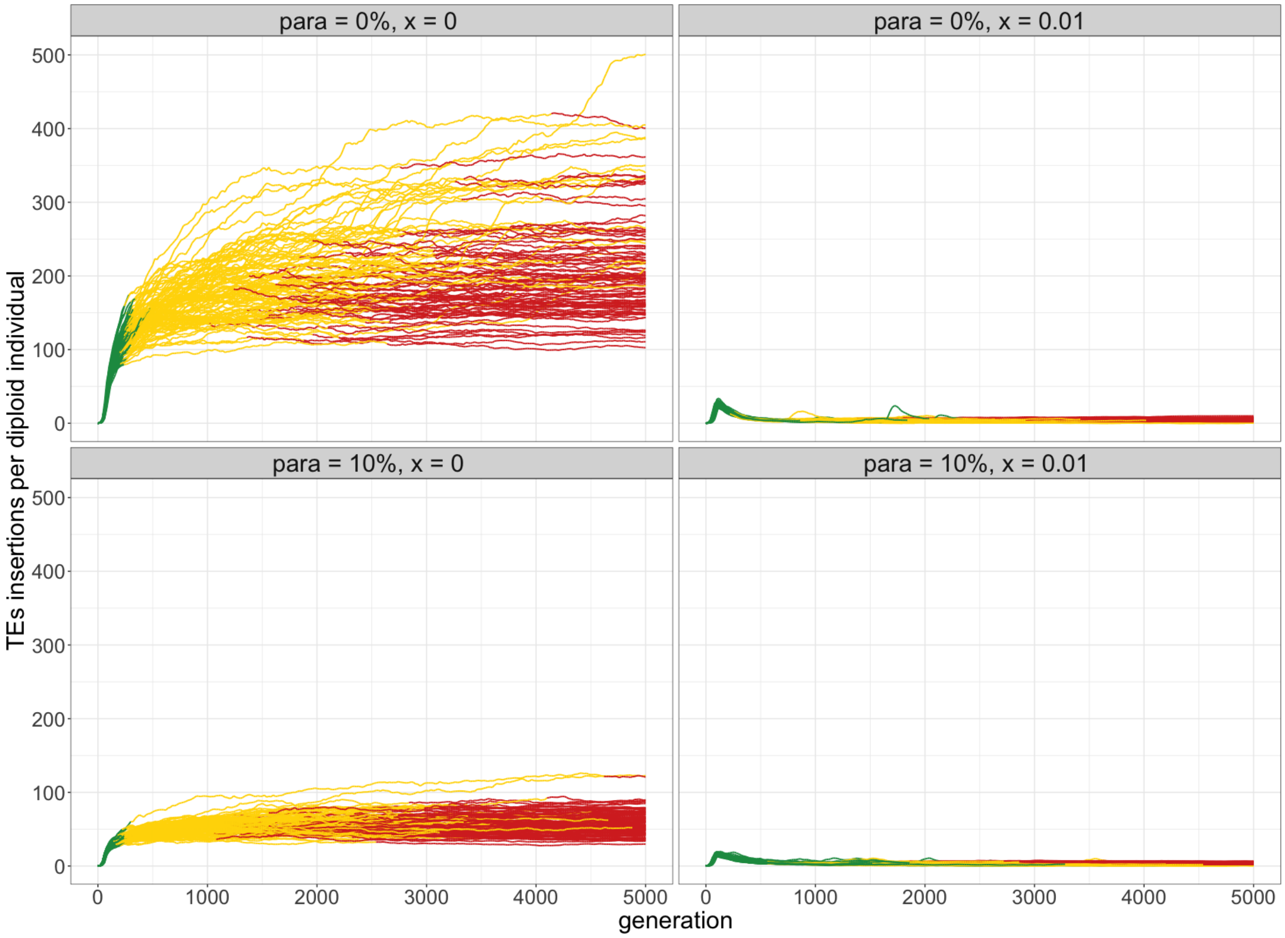


Figure 9: Invasion dynamics with neutral (left panels) and negatively (right panels) selected TE insertions. Data are shown for TE invasions with (bottom panels) and without (top panels) paramutable loci.

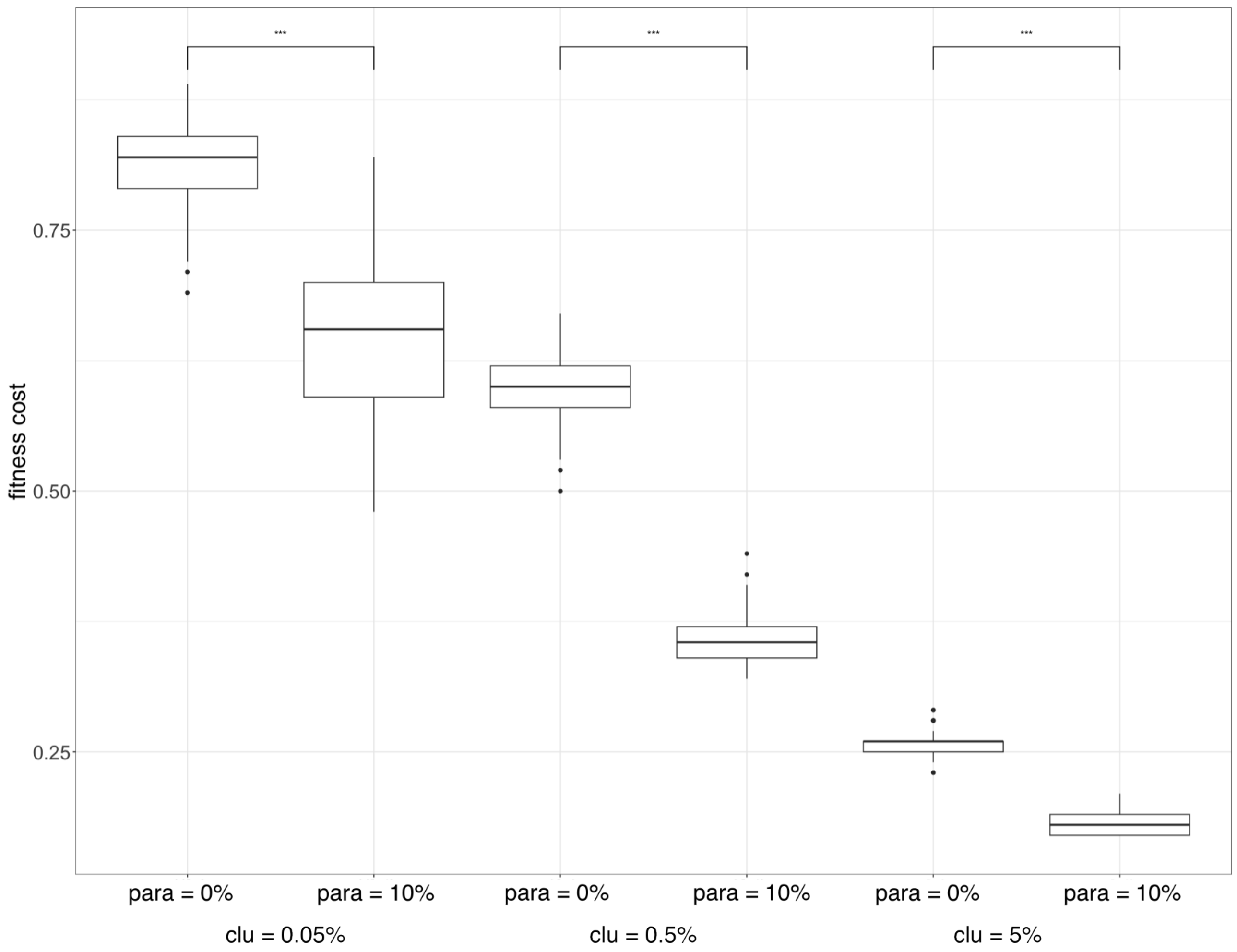


Figure 10: Paramutations reduce the fitness cost of TE invasions. We simulated TE invasions without and with 10% paramutable loci (*para*) for many different sizes of piRNA clusters (*clu*). The significance was estimated with Wilcoxon rank sum tests.

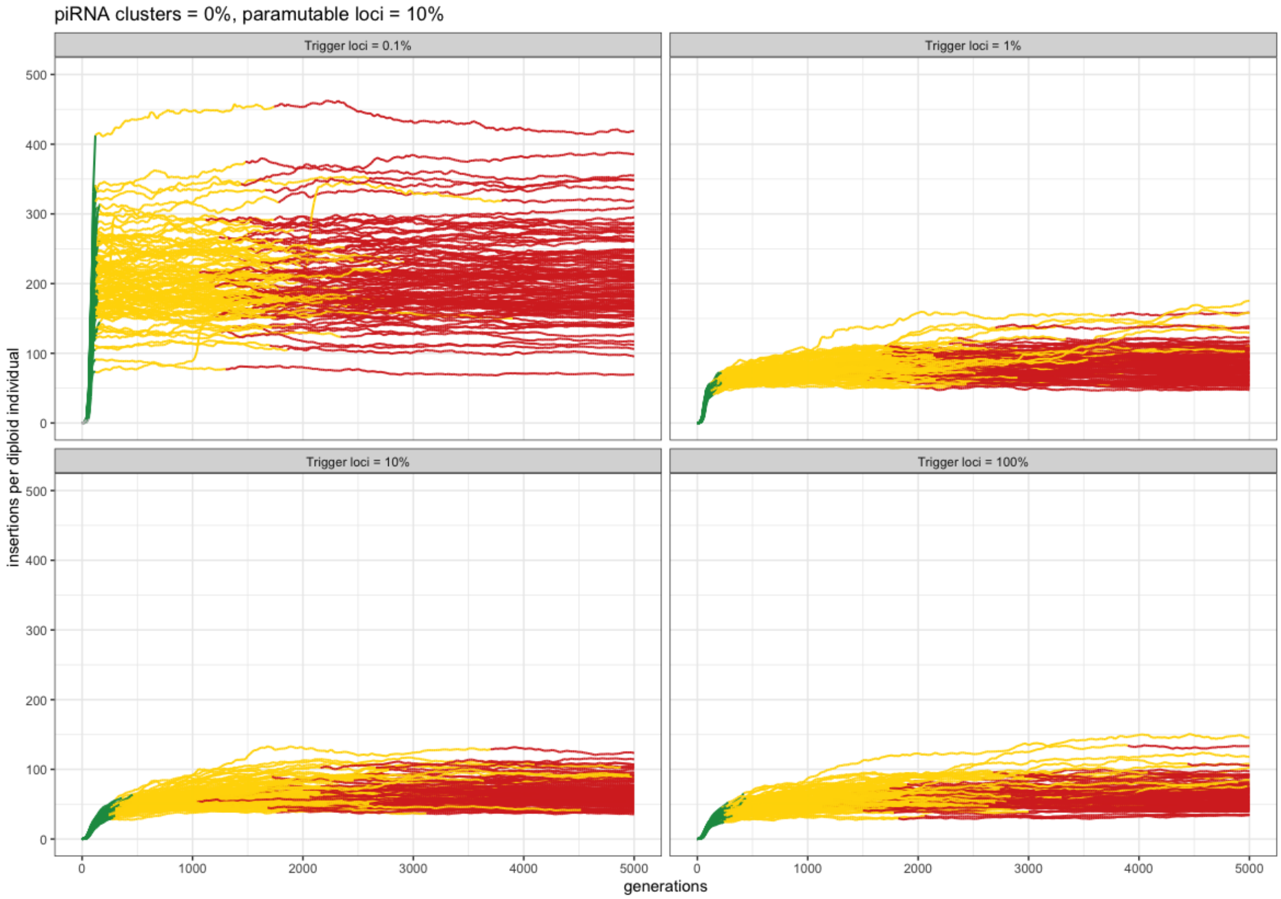


Figure 11: Dynamics of TE invasions under a siRNA model inspired by Luo et al. [2022]. We assumed that TE insertions in trigger sites (top panel) lead to the production of siRNAs that drive the conversion of TE insertions in paramutable loci into piRNA producing loci. The three phases of TE invasions can also be recognized under this siRNA model. After a rapid increase in TE copy numbers (green; rapid invasion phase), a TE invasion is initially controlled by segregating paramutated TEs (yellow; shotgun phase) and later by fixed paramutated TEs (red; inactive phase).

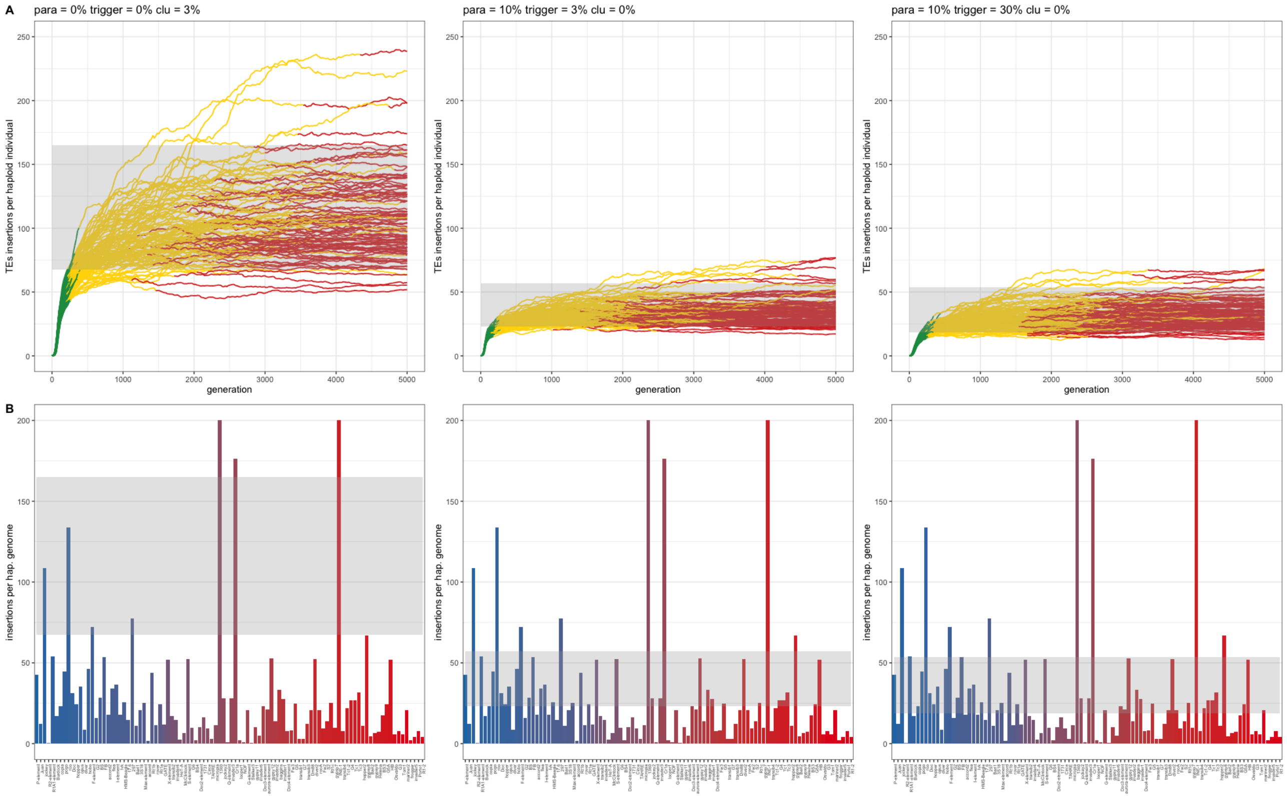


Figure 12: A siRNA model (second and third columns) predicts the observed abundance of TE families in *D. melanogaster* more accurately than the classic trap model (first column). For the siRNA model we simulated no piRNA cluster, 10% paramutable loci and 3% or 30% trigger sites. A) Abundance of TEs during simulated TE invasions. B) Observed and expected abundance of TEs in *D. melanogaster*. Bar plots indicate the abundance of each TE family. The colors of the bars indicate the average population frequency (*blue* = 0.1, *red* = 1.0). The grey shades indicate expectations from the simulations. Note that simulations with paramutable loci capture the observed TE abundance more accurately than simulations without paramutable loci. Data are from Kofler et al. [2015].

### Supplementary tables

Table 1: Fraction of stopped TE invasions after 5.000 and 10.000 generations, when insertions into piRNA clusters initiate the host response. We simulated TE invasion and counted the number of populations with a fixed piRNA producing locus (i.e. stopped invasion). Results are shown for different numbers of paramutable loci (%*para*). We used 100 replicates for each scenario. piRNA clusters had a size of 3% of the genome (%*cluster*)

| %para | %cluster | 5000 | 10000 |
|---|---|---|---|
| 0 | 3 | 0.92 | 1.00 |
| 1 | 3 | 0.97 | 1.00 |
| 10 | 3 | 0.94 | 1.00 |
| 100 | 3 | 1.00 | 1.00 |

Table 2: Fraction of stopped TE invasions after 5.000 and 10.000 generations, when insertions into siRNA-trigger-sites initiate the host response. Results are shown for different numbers of siRNA-trigger-loci (*%trigger*). We simulated 30% paramutable loci and used 100 replicates for each scenario. No piRNA clusters were simulated (*%cluster*)

| %trigger | %para | %cluster | 5000 | 10000 |
|---|---|---|---|---|
| 0 | 30 | 0 | 1.00 | 1.00 |
| 1 | 30 | 0 | 1.00 | 1.00 |
| 10 | 30 | 0 | 0.98 | 1.00 |
| 100 | 30 | 0 | 0.90 | 1.00 |
